# The nutrient sensor CRTC & Sarcalumenin / Thinman represent a new pathway in cardiac hypertrophy

**DOI:** 10.1101/2023.10.02.560407

**Authors:** Cristiana Dondi, Georg Vogler, Anjali Gupta, Stanley M. Walls, Anaïs Kervadec, Michaela R. Romero, Soda B. Diop, Jason Goode, John B. Thomas, Alexandre R. Colas, Rolf Bodmer, Marc Montminy, Karen Ocorr

## Abstract

Obesity and type 2 diabetes are at epidemic levels and a significant proportion of these patients are diagnosed with left ventricular hypertrophy. ***CREB* R**egulated **T**ranscription **C**o-activator (***CRTC***) is a key regulator of metabolism in mammalian hepatocytes, where it is activated by calcineurin (CaN) to increase expression of gluconeogenic genes. CaN is known its role in pathological cardiac hypertrophy, however, a role for CRTC in the heart has not been identified. In *Drosophila*, *CRTC* null mutants have little body fat and exhibit severe cardiac restriction, myofibrillar disorganization, cardiac fibrosis and tachycardia, all hallmarks of heart disease. Cardiac-specific knockdown of *CRTC*, or its coactivator *CREBb*, mimicked the reduced body fat and heart defects of *CRTC* null mutants. Comparative gene expression in *CRTC* loss- or gain-of-function fly hearts revealed contra-regulation of genes involved in glucose, fatty acid, and amino acid metabolism, suggesting that *CRTC* also acts as a metabolic switch in the heart. Among the contra-regulated genes with conserved CREB binding sites, we identified the fly ortholog of *Sarcalumenin,* which is a Ca^2+^-binding protein in the sarcoplasmic reticulum. Cardiac knockdown recapitulated the loss of CRTC cardiac restriction and fibrotic phenotypes, suggesting it is a downstream effector of *CRTC* we named *thinman* (*tmn*). Importantly, cardiac overexpression of either CaN or *CRTC* in flies caused hypertrophy that was reversed in a *CRTC* mutant background, suggesting CRTC mediates hypertrophy downstream of CaN, perhaps as an alternative to NFAT. CRTC novel role in the heart is likely conserved in vertebrates as knockdown in zebrafish also caused cardiac restriction, as in flies. These data suggest that CRTC is involved in myocardial cell maintenance and that CaN-CRTC- Sarcalumenin/*tmn* signaling represents a novel and conserved pathway underlying cardiac hypertrophy.

## INTRODUCTION

Despite decades of research, heart disease (HD) remains the leading cause of death in the industrialized world. An increasing number of HDs are attributed to disorders of metabolism such as obesity, insulin resistance and diabetes (1–4). Left ventricular hypertrophy has been diagnosed in as many as 50% of type 2 diabetes patients and is predictive of more adverse cardiovascular events (5). Calcineurin (CaN) has been established as part of the canonical signaling pathway underlying mammalian cardiac hypertrophy through its activation of downstream transcriptional regulators, such as Mef2 and NFAT (6–8). CaN, a Ca^2+^/CaM dependent phosphatase, also plays important roles in insulin-sensitive tissues, such as the liver, where it dephosphorylates and activates *CRTC*, a **C**REB (**c**AMP- **r**esponsive **e**lement **b**inding protein)-**R**egulated **T**ranscription **C**o-activator (9). Under basal conditions, *CRTC* is phosphorylated by SIK2, a salt inducible kinase, resulting in increased association with 14 -3-3 proteins and sequestration in the cytoplasm(10). During fasting, glucagon signaling leads to activation of cAMP-dependent protein kinase A (PKA), which phosphorylates and inhibits SIK2 while also activating *CREB*. Simultaneous activation of CRTC (by Ca2+-dependent CaN) and *CREB* (by cAMP-dependent PKA) acts as a “coincidence detector” to increase transcription of target genes involved in gluconeogenesis (9).

In mammals there are three forms of CRTC. CRTC1 is the predominant form in liver and brain, whereas CRTC2 and -3 are more ubiquitously expressed. While much is known about the role of CRTC in liver and neurons, there is little evidence to support a role for *CRTC* in cardiac function. A single study in mice, linked systemic KO of *CRTC*1 to cardiac hypertrophy, but this phenotype was likely secondary to effects on neuronal function and activation of β-adrenergic receptors (11). In addition, CRTC1 is primarily expressed in nervous tissue (12)(13) whereas CRTC2 and CRTC3 are more ubiquitously expressed (14). Here we provide evidence that *CRTC* functions autonomously in the heart. Using the fly model, we show a cardiac-specific effect of CRTC on cardiac structure and fibrosis that worsens with age. CRTC3 KD in the zebrafish model leads to similar defects in heart structure and function as in flies. We also observed functional defects in response to KD of CRTC2 & 3 in human induced cardiomyocytes. Our analysis of changes in gene expression in the fly heart in response to cardiac-specific CRTC KD and OE indicate that *Sarcalumenin* is a downstream effector of CRTC in the heart. Overall, our data suggest that *CRTC* provides an alternative pathway to NFAT in mediating cardiac hypertrophy, in part by regulating the expression of Sarcalumenin in cardiac muscle.

## RESULTS

### *CRTC* mutant flies exhibit cardiac dysfunction

Adult *Drosophila* with systemic *CRTC* knockout are sensitive to starvation and oxidative stress and are very lean (15). Neuronal rescue of *CRTC* only partially rescues these phenotypes. To determine a role for CRTC in the heart we initially used a cardiac pacing assay to examine the effects of stress on heart failure rates in *CRTC* null mutants and wildtype control flies (*w^1118^*, the genetic background from which the *CRTC* mutant line was derived (15). Male and female flies were paced with external electrodes as previously described (16, 17) using a train of square wave pulses at 6 Hz for 30 sec and hearts were scored for rhythmic beating at 60s and 120s post pacing. At 60s, approximately 40% of control hearts were beating regularly compared to less than 20% of *CRTC* mutant hearts, and by 120s post pacing, nearly 80% of hearts in control flies had recovered rhythmic function compared to only 38% of *CRTC* mutant hearts (**Supplemental Fig. 1A**). Thus, *CRTC* mutants exhibited significantly compromised heart function *in vivo*.

To examine heart function and in more detail of these *CRTC* mutants, we used denervated, semi-intact fly heart preparations (14)(18). Flies were collected upon eclosion, aged to one-week, dissected to remove the central nervous system and subsequently assayed heart function by high-speed video imaging (SOHAsoftware.com)(19, 20) Hearts from female *CRTC* mutants were significantly thinner compared to controls both during diastole (**Fig. 1A**) and systole (**Supplemental Fig. 1B**) and this cardiac restriction resulted in a significant reduction in stroke volume (**Fig. 1B**). Hearts from female *CRTC* mutants also had a significant reduction in heart period (length of one contraction cycle, **Fig. 1C**). When examining hearts from male *CRTC* mutant flies, diastolic and systolic diameters and stroke volume were reduced as in their female counterparts (**Supplemental Fig. 1D-F**), although there was no significant change in the heart period compared to controls (**Supplemental Fig. 1G**). We confirmed cardiac expression of *CRTC* with qPCR analysis of isolated hearts revealing robust expression in wildtype control flies and none in *CRTC* mutant hearts (**Supplemental Fig. 1C**).

**Figure 1.**
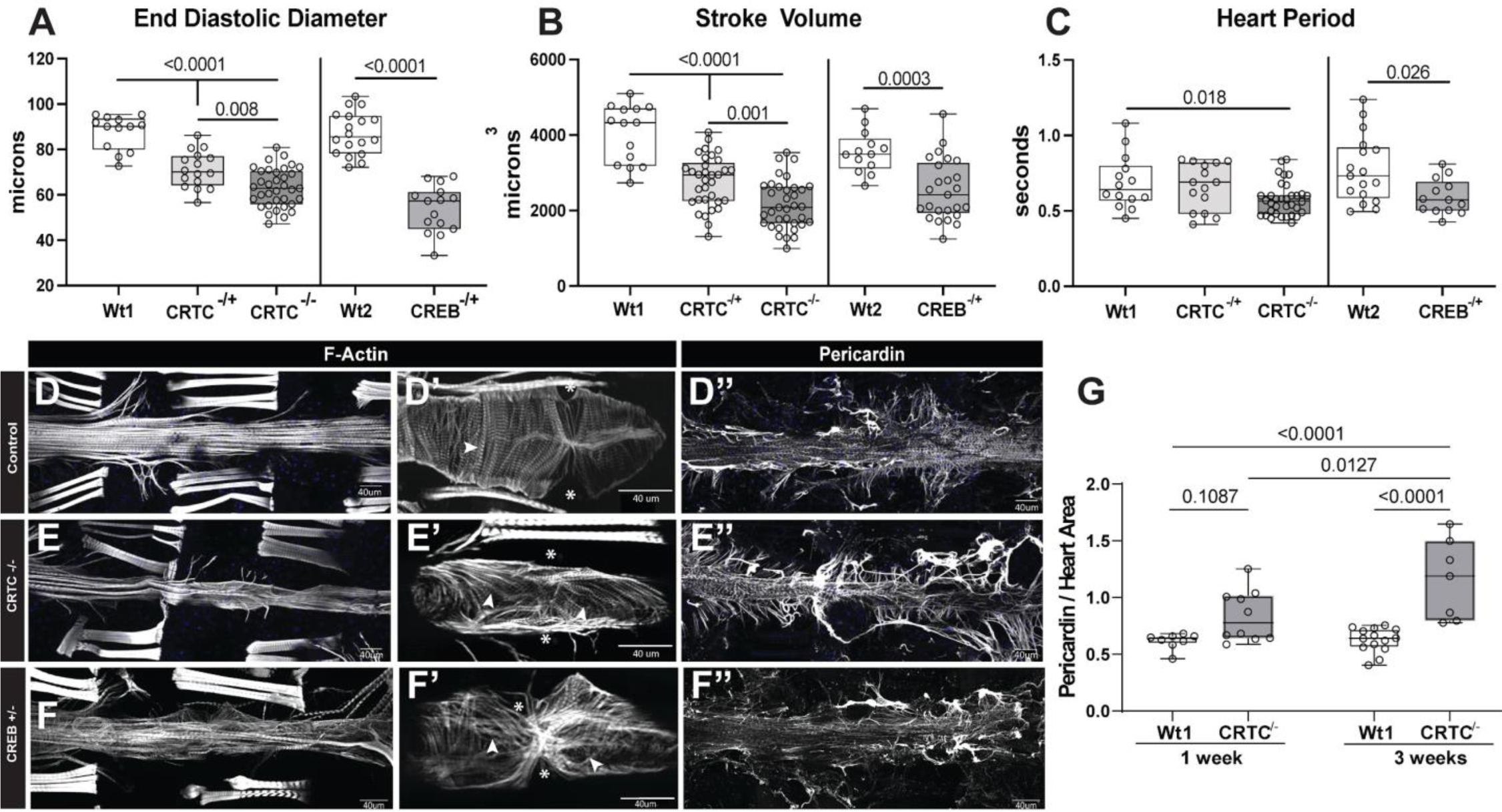
*CRTC* knockout causes cardiac dysfunction that is mimicked by *CREB* KO. **(A) Left -** Knockout of the *CRTC* gene results in significantly reduced End Diastolic Diameters (EDD) in hearts from both homozygous and heterozygous mutants compared to genetic background controls. **Right** - Similar cardiac restriction was observed in hearts from *CREB* heterozygous mutant flies compared to their genetic background controls. **(B)** Stroke volume was significantly reduced in hearts from both *CRTC-/-* and *CRTC -/+* (left) as well as in *CREB* KO flies (right). **(C)** Heart period (length of one contraction cycle) was significantly reduced in *CRTC* KO flies but not in *CRTC* heterozygotes (left). Hearts from *CREB* KO flies also showed significant reductions in HP. (For A-C all flies were 1 week old; p values determined by one-way ANOVA for *CRTC* mutants and two-tailed, unpaired student t-tests for *CREB* mutants; all plots show Max, Min and median, p values are shown). **(D)** F-actin staining with phalloidin reveals the cardiac tube and **(D’)** the tightly packed circumferential fibers (arrows) n a single chamber from a control heart (Asterisks denote position of the paired ostia, anterior is to the left in all pictures). **(E)** F-actin staining in a *CRTC* mutant exhibiting cardiac restriction and **(E’)** disorganized myofibrils (arrowheads) and ostia are malformed (asterisks). **(F)** Cardiac chamber from a *CREB* heterozygote mutant, also showing cardiac restriction, **(F’)** disorganized myofibrils and gaps (arrowheads) and malformed ostia (asterisks). **(D’’)** Wildtype control stained for Collagen IV (pericardin) reveals the extensive network of extracellular matrix that normally surrounds the heart. **(E’’)** The collagen matrix in *CRTC* and **(F’’)** *CREB* mutants is significantly expanded especially in the posterior region of the heart (to the right). **(G)** Quantification of collagen area normalized to area of cardiac actin (ImageJ) shows significant increases in both *CRTC* and *CREB* mutants compared to controls at one week (p values determined by one-way ANOVA with Tukey’s multiple comparisons post-hoc test).

The effects of *CRTC* in the liver have been shown to be mediated by interactions with *CREB* (9, 21, 22). Because homozygous *CREB* mutations are lethal we examined heart function in female CREBbD400 (15) heterozygotes (hereafter CREBb-/+) for our analysis. We observed similar reductions in heart size and stroke volume in *CREBbD400-/+* flies as seen for *CRTC* null mutants (**Fig. 1A&B**). Hearts from *CREBb*-/+ flies also exhibited significant reductions in heart period (**Fig. 1C**). Taken together these results suggested that *CRTC* loss-of-function caused heart dysfunction that likely contributed to the pacing-induced heart failure and that these effects were mimicked by reduced *CREBb* function, as previously reported for organismal metabolic effects (15).

### *CRTC* KO causes myofibrillar disorganization and fibrosis

We assessed the effects of *CRTC* loss-of-function on cardiac morphology. Following heart functional assessments, hearts were relaxed by addition of EGTA, fixed and stained for both F-actin and the cardiac extracellular matrix protein pericardin (collagen IV homolog). The *Drosophila* heart tube consists of a single layer of myocardial cells and is separated into four chambers by a series of internal valves. The myofibrils within the myocardial cells are normally arranged circumferentially ( **Fig. 1D’**, arrowhead) allowing the heart tube to contract and eject hemolymph. In *CRTC* mutant hearts, myofibrils exhibit significant disorganization with many gaps between the myofibrils (**Fig. 1E’**, arrowheads). Myocardial cells in hearts from *CREBb-/+*flies also showed disorganized myofibrils with non-circumferential orientations and gaps (**Fig. 1F’**, arrowheads). In addition, immunohistochemical staining for pericardin (collagen IV) showed that the collagen network that normally surrounds the heart tube (**Fig. 1D”)** was significantly increased around hearts of both *CRTC* mutants and *CREBb*-/+ flies compared to controls and was especially enhanced in the posterior region of the heart (**Fig. 1E”, F”).** In addition, this excess collagen deposition in CRTC mutants was further increased with age (**Fig. 1G**).

### *CRTC* mutants show normal embryonic heart development

To test whether the effects we observed in adults originated as congenital defects we examined embryonic specification of cardioblasts in stage 17 embryos. At this stage the cardioblasts are fully specified and are arranged as parallel rows of cells along the dorsal midline (**Supplemental Fig. 2A-C**). All cardioblasts express Neuromancer-1/H15 (Nmr1/H15) and in a subset that will form the ostia (inflow tracts) Seven-up (Svp), whereas the closely associated pericardial cells selectively express Zinc finger homeodomain 1 (zfh1). Both wildtype and mutant embryos stained for these markers exhibited similar staining patterns for all cell types and similar cell numbers/sizes (**Supplemental Fig. 2** compare A-C with D-F) suggesting that embryonic heart specification and development occurs normally in mutants.

### *CRTC* acts cardiac-autonomously to affect heart function

To address whether *CRTC* affects the heart cell-autonomously or systemically, we used a Gal4/UAS- mediated RNAi approach to tissue-specifically knockdown (KD) target genes. For the heart we used the myocardial cell-specific driver tinCΔ4-Gal4. TinCΔ4-Gal4 flies were crossed to UAS-CRTC RNAi lines and heart function of adult female progeny was assayed at 3 weeks of age. As for the systemic *CRTC* null mutants, cardiomyocyte-specific *CRTC* KD caused a significant reduction in heart diameters (**Fig. 2A, Supplemental Fig. 3A**) resulting in a reduced stroke volume (**Fig. 2B**), as well as reduced heart period (**Fig. 2C**). In contrast, cardiac-specific *CRTC* overexpression significantly increased diastolic, systolic diameters and heart period (**Fig. 2A&C, Supplemental Fig. 3A**) but had no effect on stroke volume (**Fig. 2B**).

**Figure 2.**
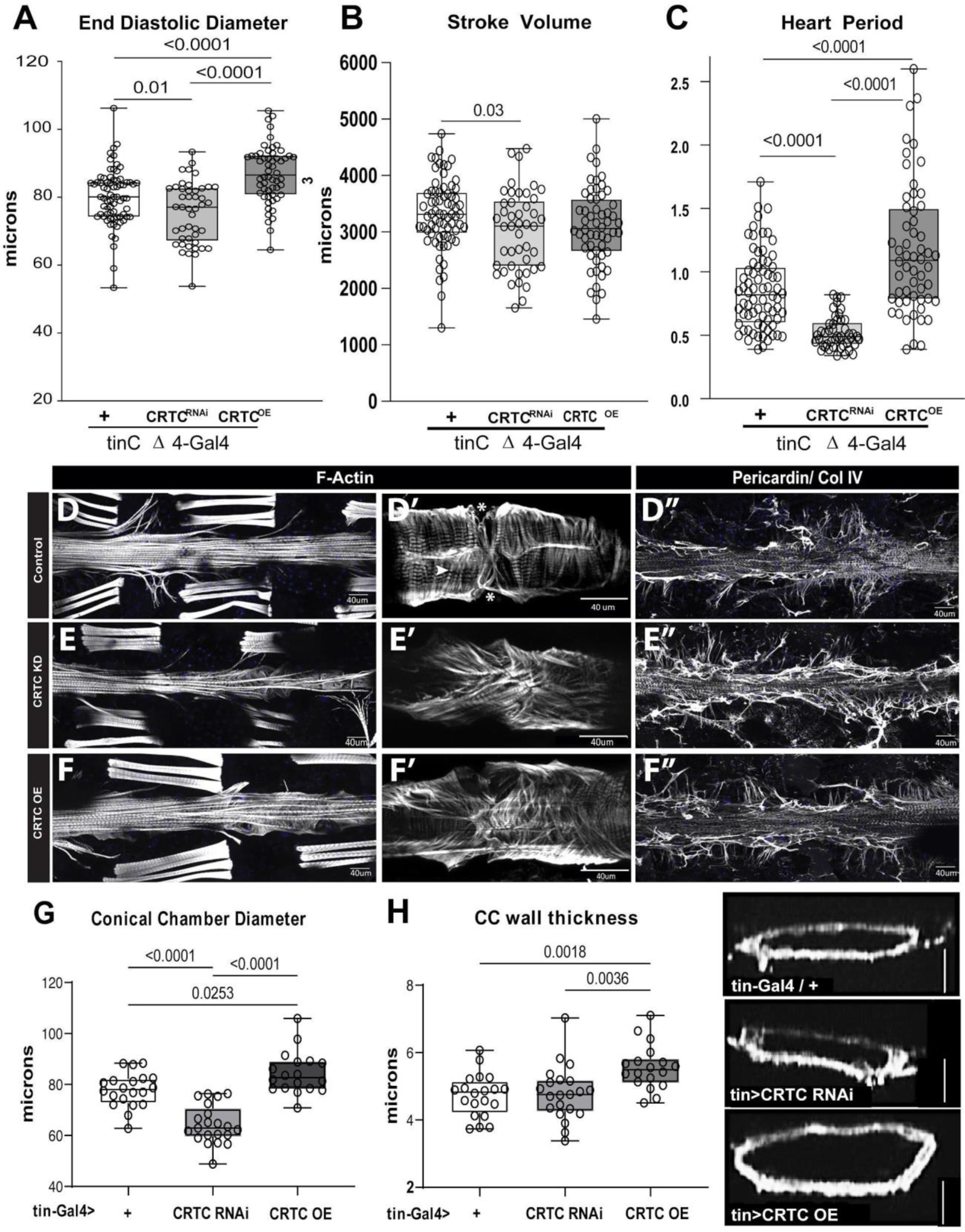
Effects of *CRTC* are cardiac autonomous. **(A)** Cardiac-specific *CRTC* KD with *tinCΔ4*-Gal4 resulted in significantly reduced EDD measured in the second cardiac chamber from movies, whereas cardiac *CRTC* OE caused an increase in EDD compared to controls. **(B)** Stroke Volume was reduced in response to cardiac *CRTC* KD. **(C)** Heart Period was significantly reduced in *CRTC* KD hearts and increased in hearts with *CRTC* OE. **(D)** F-actin staining with phalloidin reveals the cardiac tube and **(D’)** the tightly packed circumferential fibers (arrows) in a single chamber from a control heart (Asterisks denote position of the paired ostia, anterior is to the left in all pictures). **(E)** F-actin staining in a cardiac CRTC KD exhibiting cardiac restriction and **(E’)** disorganized myofibrils (arrowheads) and ostia are malformed (asterisks). **(F)** Cardiac chamber from a cardiac *CRTC* OE heart showing cardiac dilation, **(F’)** disorganized myofibrils and gaps (arrowheads) and malformed ostia (asterisks). **(G)** Diameters of dystroglycan stained cardiac conical chamber (CC) were significantly reduced with cardiac *CRTC* KD and increased with heart specific *CRTC* OE. **(H)** Cardiomyocyte thickness was measured from optical sections of dystroglycan stained hearts (shown to the right, scale bar is 50μ) and was significantly increased with cardiac CRTC OE compared to controls and cardiac *CRTC* KD hearts. All flies were 3 weeks old, plots show Max, Min and Median, significance was determined using a one-way ANOVA with Tukey’s multiple comparisons post hoc test.

To determine whether the effects of CRTC were autonomous to the cardiomyocytes or could also be achieved by manipulation of other, heart-associated cells, we examined the effects of CRTC KD in pericardial (nephrocyte-like) cells, which have previously been reported to exert effects on the heart (23). CRTC KD using the pericardial cell *Dot*-Gal4 - driver caused phenotypes opposite to those of CRTC myocardial cell KD and OE had no effect (**Supplemental Fig. 4A-C**, **Supplemental Table 1**). Since *CRTC* is prominently expressed in vertebrate neurons, we also used the pan-neuronal driver *elav*-Gal4 to modulate *CRTC* expression specifically in the nervous system (24), but observed no significant effects on cardiac function (**Supplemental Fig. 4 D-F, Supplemental Table 1**). *CRTC* also plays significant roles in liver cells by mediating the effects of glucagon signaling. In flies, fat bodies are thought to perform a number of the same functions as liver cells in humans .(25) Again, modulating *CRTC* expression using the fat body-specific driver *lsp*-Gal4 (26) did not cause any significant changes in heart function (**Supplemental Fig. 4G-I, Supplemental Table 1**). Thus, *CRTC* plays a cardiac-autonomous role in maintaining adult cardiac structure and function.

### CRTC affects somatic muscle function

Climbing assay was performed with *CRTC* systemic mutant and *CRTC* cardiac-specific KD female and male flies at 1 and 3 weeks of age. We recorded the number of flies that within 10s were able to climb above the height of 2 and 10 cm. We observed that less than 30% of *CRTC* systemic mutant flies were able to climb and cross the threshold compared to control flies where almost 80% were able to climb above the set threshold (**Supplemental Fig. 3**). This suggests that loss of *CRTC* causes muscle impairment in both females and males. Cardiac specific CRTC KD flies do not show a muscle impairment phenotype: their climbing ability is unaffected and comparable to controls. Surprisingly, cardiac specific CRTC KD female flies climb even better than controls, likely because they are overall leaner than controls and CRTC expression in their somatic muscle is unperturbed. (**Supplemental Fig. 3**).

### *Cardiac-specific manipulation of CRTC* causes cardiac remodeling, fibrosis, and hypertrophy

To examine cardiac structure more closely, fly hearts were stained for F-actin, and against Pericardin to label the collagen IV network that surrounds the heart. We found that the entire heart was restricted in response to cardiac *CRTC* KD and was enlarged in response to *CRTC* OE (**Fig. 2D-F’’**). Images of the single heart chambers at higher magnification revealed the closely packed circumferential myofibrils in control hearts as well as the significant myofibrillar disorganization in response to *CRTC* KD or OE (**Fig. 2D’-F’**). Interestingly, cardiac *CRTC* KD caused an increase in the collagen IV staining around the heart while *CRTC* OE had no effect (**Fig. 2D”-F”**). Taken together, cardiomyocyte-specific CRTC KD mimicked the effects of CRTC null mutants.

We next determined whether the enlargement we observed in response to *CRTC* overexpression was due to overall dilation of the cardiac tube or to an increase in the thickness of the myocardial cells. To measure CM thickness we focused on the conical chamber, the largest and most anterior part of the heart tube. We labeled the inner and outer plasma membrane by staining for Dystroglycan, a ubiquitous membrane bound glycoprotein. Measurements of the outer diameters from the conical chamber were similar to those obtained from brightfield images of *in situ* beating hearts, confirming the observed chamber restriction with cardiac *CRTC* KD and *CRTC* OE-dependent enlargement of the heart (**Fig. 2G**). We then quantified the thickness of the myocardial cells (**Fig. 2H, right**) using the ImageJ Trainable Weka Segmentation tool. Our results provide strong evidence that *CRTC* overexpression caused significant increases in CM thickness compared to control hearts, thus indicating hypertrophy, but cardiomyocyte thickness in KD hearts was not changed compared to controls (**Fig. 2H, left**).

### *CRTC* effects are downstream of CaN but are independent of NFAT

In the liver, *CRTC* is activated by calcineurin (CaN)-mediated dephosphorylation allowing it to enter the nucleus and modulate DNA binding by activated *CREB*. CaN OE has been reported to cause cardiac hypertrophy in fly (27) and mouse models (6). We previously provided evidence that cardiac-specific KD of the CaN fly homolog *Pp2B* caused cardiac restriction in the fly whereas overexpression of activated *Pp2B* caused cardiac enlargement consistent with CaN-mediated hypertrophy. To explore the interactions between *CRTC* and *CaN* in the fly heart further, we overexpressed constitutively activated Pp2B in *CRTC* mutant flies. We observed that in 3 week old flies, cardiac OE of activated Pp2B/CaN caused a significant enlargement of the heart and increased heart period (bradycardia) (**Fig. 3A&B**). These effects were similar to those seen in response to cardiac *CRTC* OE (**Fig. 2A,C**). Importantly, the hypertrophy in response to Pp2B/CaN OE was observed to be reduced by reductions in *CRTC* expression in a dose-dependent manner (**Fig. 3A&B**), suggesting that *CRTC* mediates some of the downstream effects of CaN in the heart.

**Figure 3.**
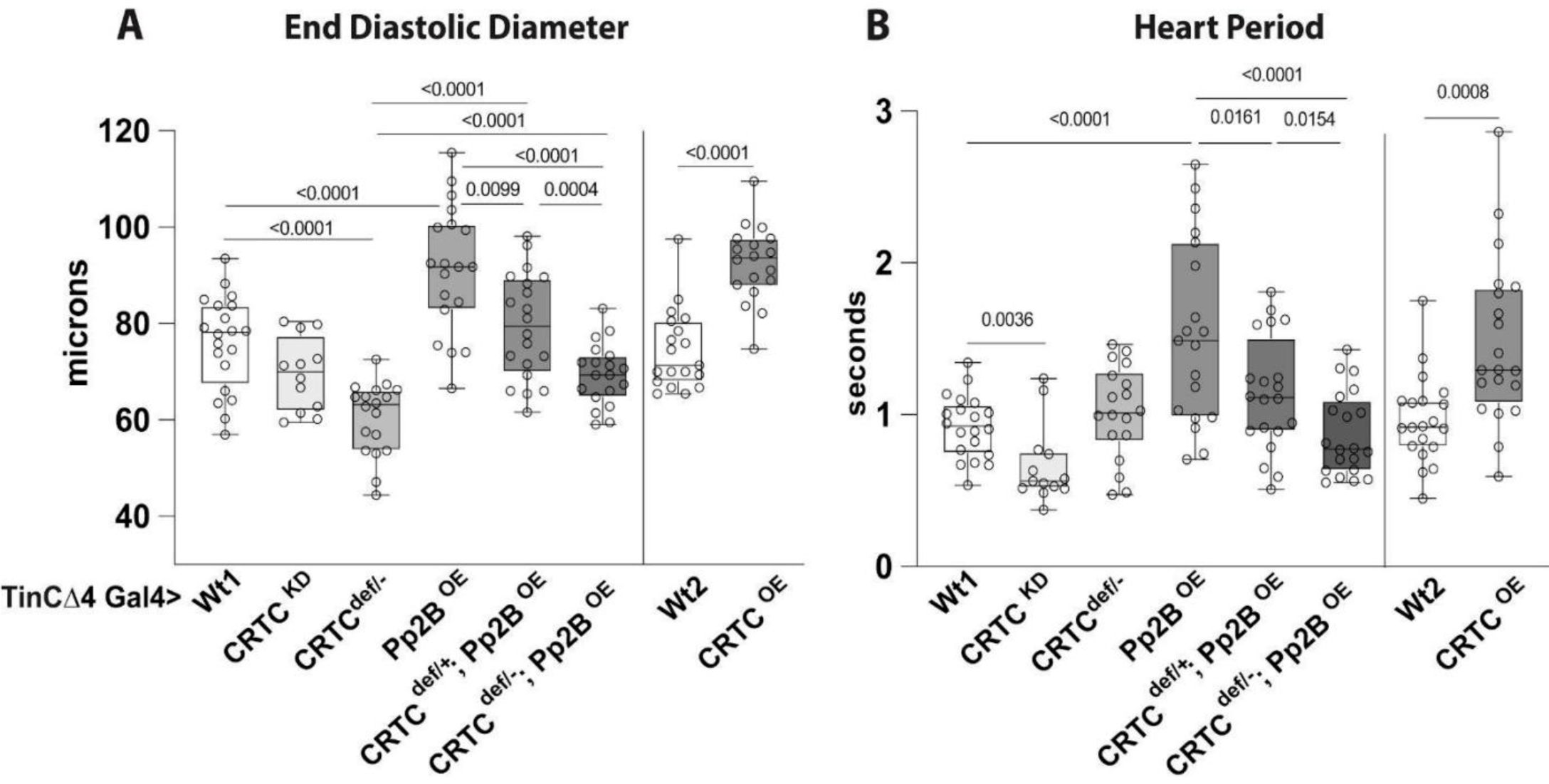
Hypertrophic effects of Calcineurin / Pp2B are mediated by *CRTC* in the heart. **(A) Left –** End Diastolic Diameters are reduced in cardiac- specific *CRTC* KD and mutant (*CRTCΔ/-)* compared to their genetic background controls. Constitutive cardiac over-expression of *Pp2B* caused an increase in EDD that was progressively rescued in *CRTC* heterozygous and homozygous mutant backgrounds. **Right -** Cardiac- specific *CRTC* OE caused a significant hypertrophic response similar to cardiac *Pp2B* OE. **(B) Left -** Cardiac-specific *CRTC* KD caused tachycardia (shorter heart periods) compared to genetic background controls. *Pp2B* OE caused bradycardia (longer heart periods) that was progressively rescued by reducing CRTC expression. **Right -** *CRTC* OE also caused (bradycardia) compared to genetic background controls. Plots show Max, Min and Median, significance was determined using a one-way ANOVA with Tukey’s multiple comparisons post hoc test.

In vertebrates, the effects of CaN OE on cardiac hypertrophy were reported to be mediated by the transcription factor NFAT (27)(28). We used tinCΔ4-Gal4 driver to heart-specifically KD to test a cardiac role for NFAT and we also tested for interactions with CRTC in a “sensitized” heterozygote background. Cardiac KD of NFAT alone had no effect on heart size and there was no genetic interaction between CRTC+/- and NFAT (**Supplemental Figure 5**). These results suggest that NFAT does not affect heart structure and function in flies, nor does it act through CRTC.

### *CRTC* is highly expressed in the zebrafish heart and KD causes cardiac restriction

We used zebrafish to examine the role of *CRTC* in a vertebrate model. Hearts were isolated from adult fish and pooled (10 month old fish, 8 hearts per sample). qPCR analysis suggests that all three forms of *CRTC* were expressed in the heart, but *CRTC3* was the most prominent form (**Supplemental Fig. 6A).** We injected morpholinos targeting *CRTC*3 into fertilized embryos at the one cell stage and analyzed heart function at 72 hours post fertilization (hpf), a stage when the fish are still transparent and the heart can be readily visualized. *CRTC*3 KD did not noticeably affect the overall morphology of the fish or on tail musculature and fin development was unperturbed (**Fig. 4C&D**). Analysis of heart function indicated that the primary effect of *CRTC* 3 KD was a significant reduction in the diastolic and systolic surface areas of KD fish compared to controls (**Fig. 4A&B**). This cardiac restriction occurred in both atria and ventricles but was most prominent in the ventricles and paralleled the heart diameter restriction seen with *CRTC* KD in the fly heart (**Fig. 1**).

**Figure 4.**
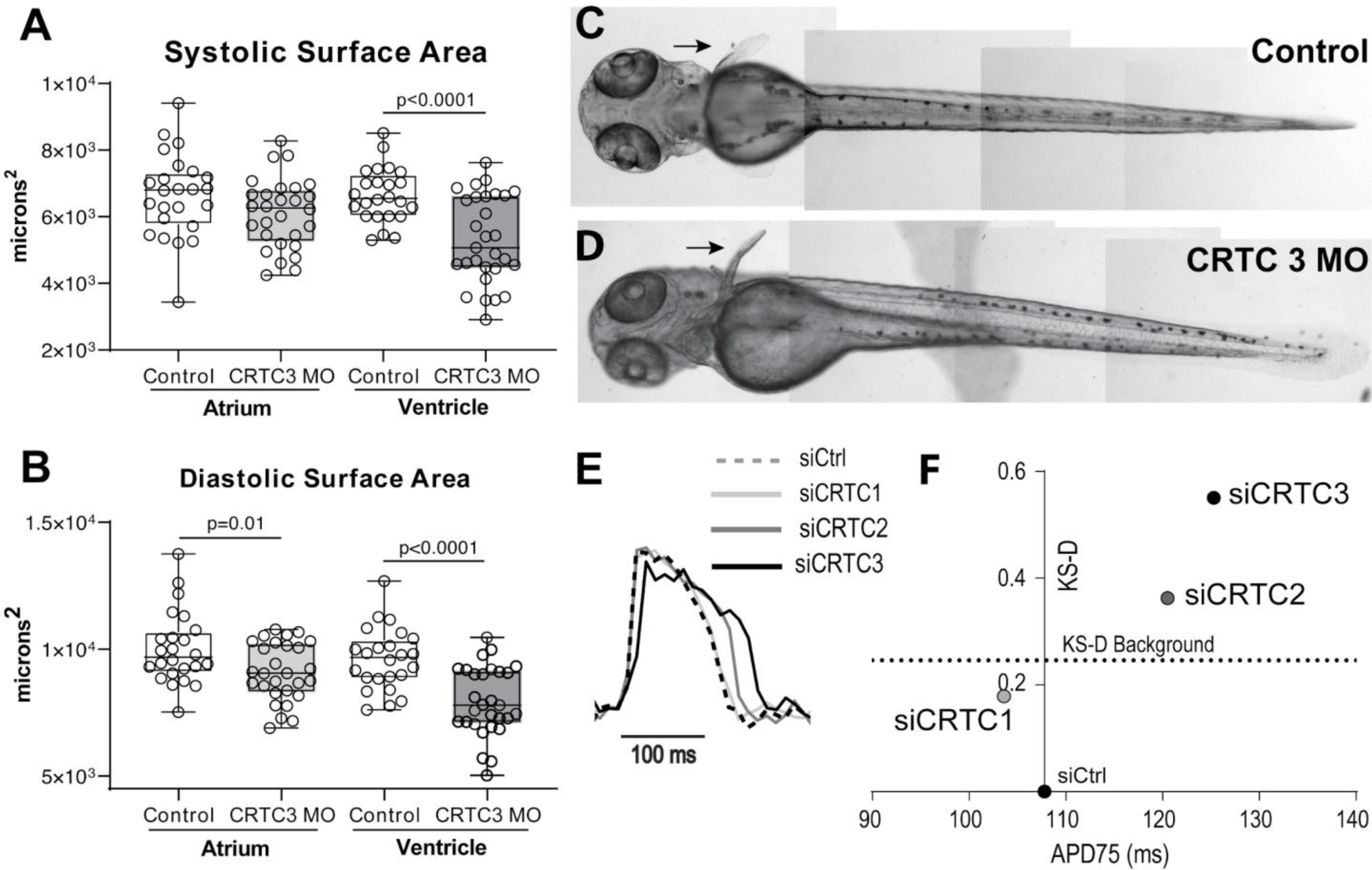
*CRTC* affects cardiac function in zebrafish and APD in hiPSC-Cardiomyocytes. **(A)** End Diastolic chamber size and **(B)** End Systolic chamber size in atria (left) and ventricles (right) of 72 hpf fish show significant reductions in surface area, especially in the ventricles, following MO KD of *CRTC 3*. Fish were 72 hpf; plots show Max, Min and median, significance was determined using Student’s unpaired T-test, p values are shown. **(C)** Images of control and **(D)** *CRTC* KD fish show normal development of body wall muscle and fins (arrows). **(E)** Action Potentials in cardiomyocytes, which were induced to differentiate from hiPSCs on Day 0. Cells were transfected with siRNAs against all three *CRTC*s on Day 25 and on day 28 were incubated with Fluo Volt and subjected to kinetic imaging to obtain cell specific voltage traces. Representative voltage traces for control siRNA and *siCRTC* transfected cells. **(F)** K-S distance plot of APD75 versus K-S distance shows significant increases with KD of either *CRTC 2* and *CRTC3* but not KD of *CRTC1*.

### *CRTC* is localized to Z bands in fish and fly hearts

Fly hearts were also stained with CRTC antibodies (gift from Marc Montminy) to localize CRTC protein. As expected, we observed prominent CRTC staining in myocardial cell nuclei ( **Fig. 5A&B**). In addition, we also saw discrete bands of CRTC staining associated with bands of α-actinin staining (**Fig. 5**) in both in the overlying, non-cardiac longitudinal fibers (**Fig. 5 C&D**) and in the myocardial cells (**Fig. E-G’**) and suggesting that *CRTC* is localized to Z bands in muscle. To confirm that the antibody used in the CRTC staining was specific to CRTC we also examined hearts in a fly line with Flag-tagged CRTC. Hearts were probed with anti-flag antibodies and we observed similar staining patterns as for the CRTC antibody (**Supplemental Fig. 7 C&D**).

**Figure 5.**
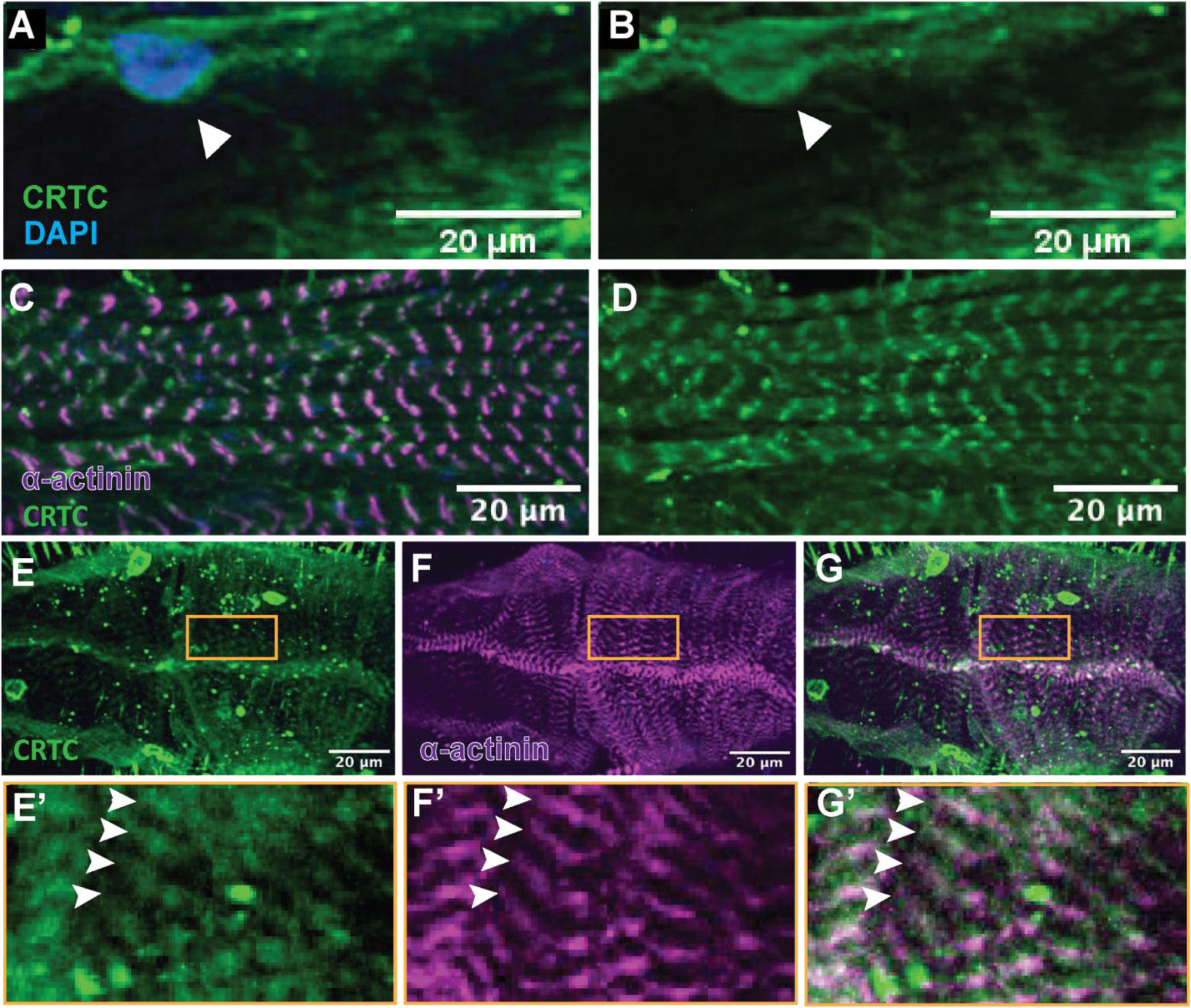
CRTC localizes to nuclei and Z bands in fly hearts. **(A-B)** Nuclear CRTC localization in cardiomyocytes (arrowheads); CRTC staining is green, nuclear staining (blue) is with DAPI. Scale bars 20 µm **(C-D)** Co-staining of non-myocardial, ventral longitudinal fibers with CRTC and α-actinin antibodies show CRTC strongly associated with the Z-bands. **(E- G)** Optical section through the chamber of the heart tube stained for CRTC (E, green), α-actinin (F, magenta), and the merged image (G). Scale bars are 20 µm. **(E’-G’)** Higher magnification of regions in yellow rectangles. Arrowheads show location of anti-CRTC staining showing co-localization with the Z- bands (white).

We also co-stained larval zebrafish hearts with antibodies against *CRTC* and MF20 (myosin). We observed *CRTC* localization in both nuclei (**Supplemental Fig. 7A&B**) and also associated with regions between the M-bands, the region where Z bands are located (**Supplemental Fig. 7 B’&B”**).

### RNA Seq analysis of *CRTC* KD and OE hearts reveals concerted regulation of cardiac metabolic pathways

To examine the effects of CRTC on cardiac gene expression we performed RNAseq analysis of isolated fly hearts. Hearts from cardiomyocyte-specific (tinCΔ4-Gal4) *CRTC* KD flies exhibited upregulation of 357 and downregulation of 186 transcripts whereas cardiac CRTC OE resulted in the upregulation of 436 and down regulation of 392 transcripts relative to their respective control flies (Fold change ±1.25 Fold; adj. q-value <0.05) (**Fig. 6A**). Comparative analysis revealed that a total of 140 genes were contra-regulated between CRTC RNAi and OE flies, suggesting CRTC-dependent regulation of specific transcriptional programs (**Fig. 6A, Supplemental Table 5, Supplemental Fig. 8**). Gene Ontology (GO) analysis shows that loss of CRTC enhanced, while gain of *CRTC* suppressed, oxidative-phosphorylation, innate immune response glycolysis/gluconeogenesis, amino acid biosynthesis and fatty acid elongation programs. Conversely, loss of *CRTC* suppressed and gain of *CRTC* enhanced expression of fatty acid degradative genes (**Fig. 6B**).

**Figure 6.**
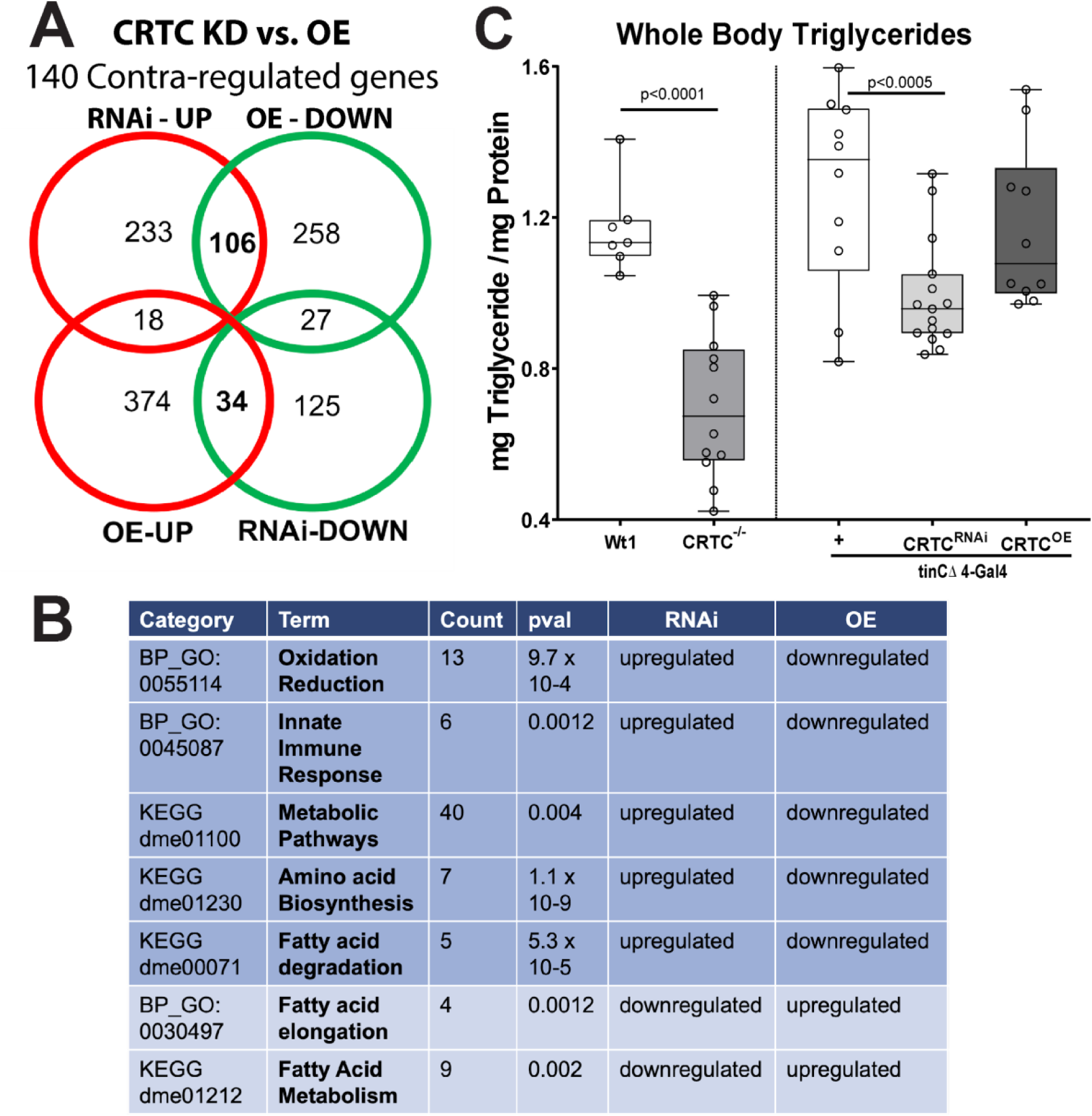
Cardiac *CRTC* KD and OE concertedly regulate metabolism in the heart. **(A)** Venn diagram showing up and down regulated genes in response to cardiac-specific KD or OE of *CRTC*. Bolded numbers in the center represent genes that are concertedly regulated by CRTC; 106 are DOWN with *CRTC* OE and UP with *CRTC* KD, 34 are UP with *CRTC* OE and DOWN with *CRTC* KD. **(B)** Significantly affected GO categories for the 140 concertedly regulated genes are primarily pathways involved in metabolic regulation. **(C)** Whole body triglyceride levels are significantly reduced in *CRTC* systemic mutant flies (left) and are also significantly reduced in cardiac-specific CRTC KD flies. Plots show Max, Min and Median, significance was determined using an unpaired T-test (C left) and a one- way ANOVA with Tukey’s multiple comparisons post hoc test (C right); p values are shown.

These results are consistent with previous observations that *CRTC* null fly mutants exhibited dramatically reduced lipid stores, a reduction that could only be partially restored with neuronal OE of *CRTC* (*15*). Because our RNAseq results suggest that *CRTC* regulates metabolism in myocardial cells and because the heart is an energetically demanding organ, we wondered if cardiac *CRTC* levels also played a role in systemic lipid storage. We analyzed whole body triglyceride content in CRTC null mutants as well as with cardiac-specific modulation of *CRTC* expression. As previously reported, we found that CRTC mutants exhibited a dramatic reduction in systemic triglyceride levels (**Fig 7C**, left). Remarkably, cardiac-specific KD of *CRTC* also resulted in a significant decrease in the whole-body fat levels (**Fig. 7C**, right). Taken together, these data suggest that CRTC may act as a metabolic switch in the heart to regulate lipid, carbohydrate and protein metabolism. Interestingly, CRTC KD in the heart seems to have a systemic effect on organismal metabolism, potentially as a secreted cardiokine (see discussion).

**Figure 7.**
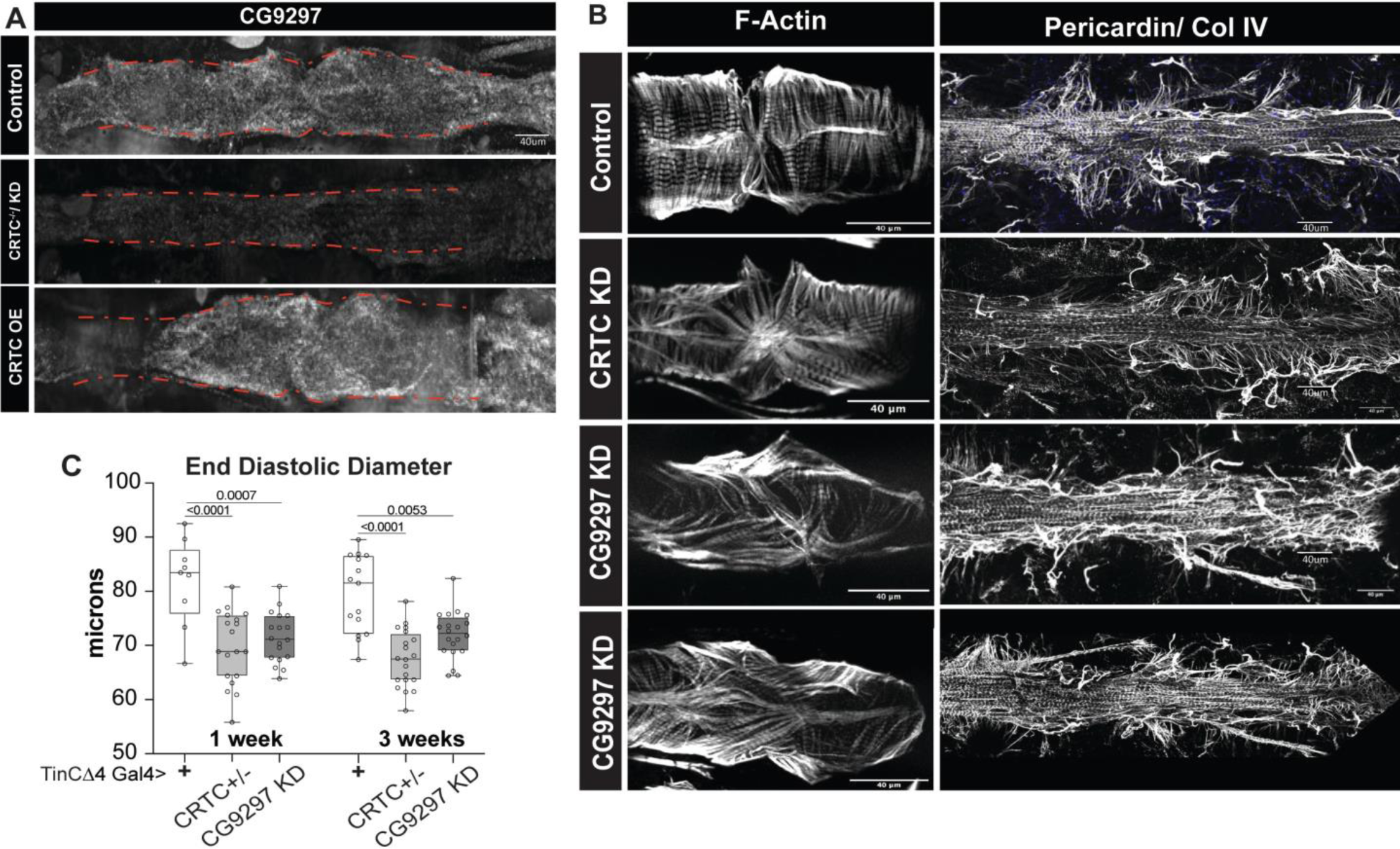
*CG9297 / Sarcalumenin* is regulated by *CRTC* in the heart. **(A)** HCR for *CG9297* in the Drosophila hearts shows that *CG9297* is ubiquitously expressed in hearts from control flies, expression is significantly reduced in *CRTC* mutant and *CRTC* KD hearts and increased in *CRTC* OE hearts. **(C)** F-actin staining shows disorganized myofibrils in CRTC and CG9297 KD compared to the closely packed myofibrils in control. Anti-Pericardin (collagen IV) staining shows a more extensive collagen network surrounding the *CRTC and CG9297* KD heart compared to control. **(D)** End diastolic diameters are significantly reduced in both CRTC heterozygotes and in response to cardiac CG9297 KD at 1- and 3-weeks of age compared to wild-type controls (+).

### *Sarcalumenin* / *thinman (tmn*) is a candidate downstream effector of *CRTC*

We next wanted to identify potentially direct targets of CREB/CRTC among the 140 contra-regulated candidate genes from our RNASeq data. We first determined which Drosophila genes contained a *CrebA* or *CrebBb* binding site around the transcriptional start sites (-2kb upstream/ +1kb downstream). We used binding site motifs from FlyFactorSurvey (29) and genes were chosen only if the binding sites were conserved between two divergent fly species, *Drosophila melanogaster* and *Drosophila persimilis.* This approach yielded 915 unique genes (309 with CrebB sites, 714 with CrebA sites, 108 with both; **Supplemental Table 2**). We tested if this approach allowed for enrichment of direct CRTC/CREB targets. Indeed, genes with CREB binding sites in the dysregulated set of genes were more abundant (12%) compared to all genes (9.2%) or dysregulated genes with no CREB binding sites (8.9%) (hypergeometric test; p-value < 0.05, **Supplemental Table 3**). The 140 contra-regulated candidate genes from our RNASeq data (**Supplemental Table 4)** were subsequently filtered for genes that were expressed in myocardial cells as determined by single cell sequencing analysis (30). We identified 15 genes that met all these criteria (**Supplemental Table 5**). Among them, *CG9297* was upregulated upon *CRTC* OE and downregulated with CRTC RNAi; it had the strongest cardiac expression of genes with human orthologs and had the consensus CREB binding site sequence TA<u>TGACGT</u>GGCT (half-site). *CG9297* is orthologous to the human gene *Sarcalumenin*, a Ca^2+^ binding protein that buffers and transports Ca^2+^ within the sarcoplasmic reticulum (SR) in skeletal and myocardial cells (31).

To confirm CRTC regulation of expression, we performed *in situ* hybridization chain reaction (HCR) on exposed fly hearts and found that *CG9297* mRNA is expressed throughout the *Drosophila* cardiac tube (**Fig. 7A**). Expression of both *CRTC* and *CG9297* was significantly reduced in hearts from *CRTC* mutants as well as in hearts with cardiac specific *CRTC* KD (**Fig. 7A**). Conversely, hearts with cardiac-specific *CRTC* OE showed increased *CG9297* mRNA expression (**Fig. 7A**). These results are consistent with our RNAseq data and suggest that CRTC plays a role in regulating *CG9297* expression. We further examined the effects of *CG9297* KD on hearts in our fly model. RNAi-mediated, cardiomyocyte-specific KD of *CG9297* caused cardiac restriction and myofibrillar disarray already in one week old flies (**Fig. 7B&C**) similar to the effects of *CRTC* KD and CRTC null mutant flies and CRTC KD in fish (**Fig. 1&4**). In addition, we observed a fibrotic phenotype (**Fig.7B**) similar to *CRTC* null mutant and *CRTC* KD fly hearts (**Supplemental Fig. 6C-E’’**). Due to this restricted heart phenotype we propose to rename *CG9297* as *thinman* or *tmn*. Further, functional analysis of flies with cardiac *tmn* KD also showed a progressive, age-dependent prolongation of systolic intervals (SI) in response to cardiac *tmn* KD (**Fig. 8A**). We see similar age-dependent increases in SI with cardiomyocyte *CRTC* KD (**Fig. 8B**). We have previously shown that SI tracks with intracellularly recorded action potentials (AP) and can be used as a surrogate for AP duration (APD) in this single cell layer heart tube (34). Reductions in *Tmn/SRL* expression would be expected to cause decreased SR Ca^2+^ buffering and an increase in cytosolic Ca^2+^ levels (see **Model Fig.8**) resulting in prolonged APD/SI.

**Figure 8.**
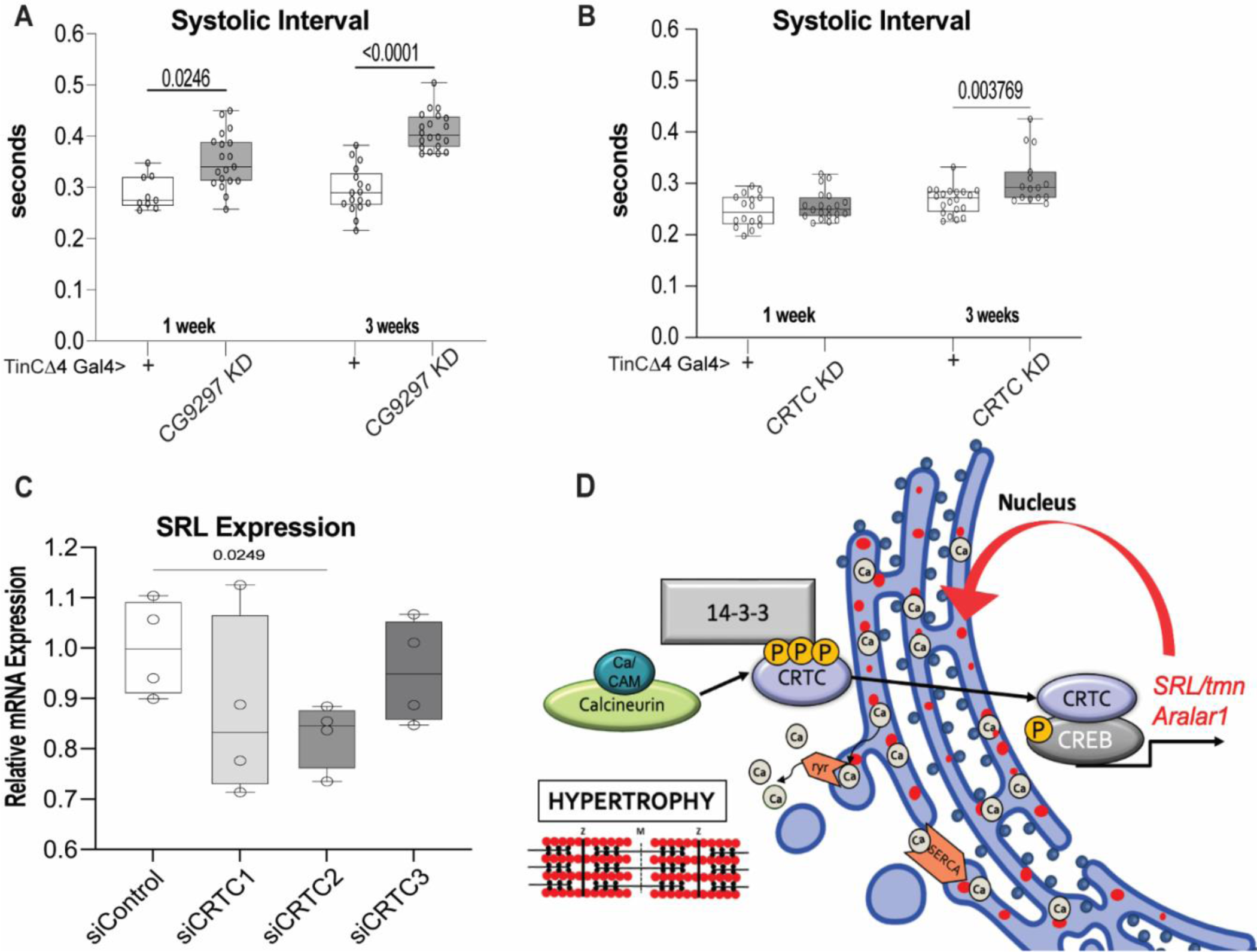
Model of CN - CRTC - *SRL* signaling pathway. **(A)** Systolic interval is significantly increased in response to cardiomyocyte KD of CG9297 / thinman at 1- and 3-weeks of age compared to wild-type controls (+). **(B)** Systolic interval is significantly increased in response to cardiomyocyte KD of CRTC at 1- and 3-weeks of age compared to wild-type controls (+). **(C)** RT-qPCR data showing *SRL* mRNA expression in IPSC-CMs after CRTC KD (significant after CRTC2 KD). Plots show Max, Min and Median, significance was determined by one-way ANOVA with Tukey’s multiple comparisons post hoc test, p values shown. **(D)** Model of CaN-CRTC-*SRL* signaling.

### *CRTC* KD affects action potential duration in hiPSC-cardiomyocytes

Finally, we used human induced pluripotent stem cells that had been induced to form cardiomyocytes (hiPSC-CMs) to determine if *CRTC* plays a role in human cardiomyocyte function. We induced CM differentiation (32, 33) and at day 25 of differentiation, cells were dissociated, replated and transfected with siRNAs against all three isoforms (**Supplemental Table 6**). Three days post transfection, cells were loaded with a voltage sensitive dye, imaged and processed generating single cell voltage traces for all cells in each well. Representative traces of median APs from cells with KD of *CRTC*1,2&3 compared to controls are shown in **Fig. 4E**. Action potential duration (APD) was quantified from individual cells at 50, 75 and 90% repolarization and showed that KD of *CRTC* 2 and *CRTC*3 significantly increased the APD at all points during repolarization (**Fig. 4 E,F, Supplemental Fig. 6B-H**). However, KD of *CRTC* 1, the primary form expressed in neurons, did not significantly affect APD duration relative to controls (**Fig. 4 E,F**).

In light of our results in the fly heart, we quantified *SRL* expression in these hiPSC-CMs in response to CRTC KD. We observed significantly reduced expression of *SRL* in response to KD of *CRTC2* (**Fig. 8C**). These data suggest CRTC 2 and/or 3 have specific roles in vertebrate myocardial cell function that are distinct from that of CRTC1. Reduced expression of *SRL* would be expected to cause increases cytosolic Ca^2+^ levels with associated increases in APD. Finally, APD prolongation in hiPSC-CMs is consistent with the observed increases in SI in the fly heart in response to cardiac *CRTC* and *SRL* KD (**Fig. 98**).

## DISCUSSION

Obesity, diabetes and metabolic syndrome are prominent risk factors thought to contribute to a number of cardiomyopathies. ***CRTC***, a ***CREB* R**egulated **T**ranscription **C**o-activator, is known to be a key regulator of metabolism in mammalian hepatocytes and neurons, but despite its prevalent expression in the heart, a cardiac-autonomous, regulatory role for *CRTC* has yet to be identified. Our data here show that hearts from *CRTC* mutant flies exhibit significant cardiac restriction, myofibrillar disorganization and fibrosis (**Fig. 1**). Using multiple tissue-specific drivers to KD *CRTC* we found that only cardiac KD recapitulates the restriction, myofibrillar disorganization, and fibrosis seen in the mutants (**Fig. 2, Supplemental Fig. 4**). Cardiac KD of either the *CRTC* co-modulator *CREBb* or the *CRTC* activator calcineurin (CaN, fly homolog Pp2B) recapitulates the *CRTC* mutant phenotype (**Fig. 3**). Finally, the cardiac-specific effects of *CRTC* appear to regulate the maintenance of adult myocardial cells, as heart development was seemingly unaffected in the *CRTC* systemic mutants (**Supplemental Fig. 2**).

Our data from zebrafish suggest that a cardiac role for *CRTC* is conserved. All three isoforms of *CRTC* (1,2,3) were detected in isolated hearts from adult fish, but *CRTC* 3 had by far the highest expression levels (**Supplemental Fig. 6A**) and KD of *CRTC* 3 caused significant cardiac restriction, similar to the fly cardiac phenotype (**Fig. 4 A,B**). In addition, all three *CRTC*s were also expressed in 25 day hIPSC-cardiomyocytes (iPSC-CMs), but *CRTC1* was expressed at very low levels compared to *CRTC 2 & 3,* which were expressed at levels 5-6 times higher (**Supplemental Table 7** ) (34). Consistent with this, siRNA KD of *CRTC* 2 or 3 in hiPSC-CMs caused significant action potential broadening while KD of the predominantly neuronal form, *CRTC*1, had no effect (**Fig. 4 E&F, Supplemental Fig. 6**). Also consistent with these results is a report that systemic KO of *CRTC*1 had no effect on cultured mouse cardiomyocytes (11). On the other hand, this same study documented increased ventricular *CRTC*1 levels in patients with aortic stenosis and hypertrophic cardiomyopathy, however the authors did not analyze the *CRTC* 2 or 3 isoforms, which are the predominant cardiac isoforms (14).

One way that *CRTC* may regulate myocardial cell function is by controlling metabolic pathways in the heart as it does in mammalian liver (9). CRTC null mutants, as previously reported (35), or cardiac KD of CRTC (this report) exhibit a lean phenotype, *CRTC* null mutant flies had significantly reduced body TAG levels. In a surprising result, flies with cardiac-specific *CRTC* KD also had significantly reduced whole body fat. This may account for the observation that neuronal *CRTC* OE did not completely rescue TAG levels in previous studies (15). Comparisons of RNA seq data between hearts with cardiac *CRTC* KD and *CRTC* OE suggested that *CRTC* acts as a metabolic switch between fatty acid metabolism and synthesis in the heart. The fly data therefore suggests cardiac metabolism impacts systemic fat content. A similar situation was observed in mice and flies, where cardiac KD of a subunit of the mediator complex (Med13) makes the animals prone to fat accumulation in response to a HFD, whereas cardiac OE protects them from the effects of HFD (36, 37). In flies but cardiac OE of CRTC had no significant effect on systemic fat accumulation. It will be interesting to see if the systemic leanness of cardiac KD flies also depends on Med13/skd function.

CRTC has been implicated in skeletal muscle hypertrophy (reviewed in (38) and our data from flies also suggest a separate role for *CRTC* in somatic muscle function, assayed by climbing ability. In a previous study we reported no effect on climbing in *CRTC* mutants, but in that study only very young flies were tested (3-5 days old) (35). We tested older flies (1 and 3 weeks) and found that climbing was impaired in systemic *CRTC* mutants and that performance in both wt and mutant flies declined in an age-dependent manner. Interestingly, flies with cardiac-specific *CRTC* KD performed the same as or slightly better than controls (**Supplemental Fig. 3C,E**) suggesting that somatic muscle function was compromised in the systemic *CRTC* mutants but not in the cardiac KD flies, despite the systemic effect of cardiac CRTC KD on organismal fat content.

Overexpression of CaN has been shown to induce cardiac hypertrophy in mice and flies (6–8) (25) and we again observed that OE of CaN/Pp2B in the fly heart caused hypertrophy (**Fig. 2**). This hypertrophy could be partially reversed in *CRTC* heterozygotes and was further reduced in *CRTC* homozygous null mutants (**Fig. 3**). These data all point to a cardiac-autonomous role for *CRTC* acting downstream of CaN in the fly heart. Further, the lack of a robust effect of NFAT KD on heart function in the fly (**Supplemental Fig. 5**) suggests that *CRTC* signaling may mediate a novel hypertrophic signaling pathway. CRTC has been shown to modulate the activity of other transcription factors such as AP1 and ATF6, as well as non-canonical roles, for example preventing SREBP activation by sequestering Sec31 in the cytosol and possibly even as a secreted signal (reviewed in (21). Thus, CRTC may play roles in cardiac function beyond the regulation of gene expression.

We identified a number of genes that were contra-regulated in response to *CRTC* KD and OE, that are expressed in myocardial cells and have upstream *CREBb* binding sites conserved between distant *Drosophila* species. Our examination of one of these genes, *thinman (tmn),* orthologous to vertebrate *Sarcalumenin*, had cardiac-specific effects that paralleled those seen with *CRTC* KD and KO (**Fig. 6,7, Supplemental Fig. 8**). Sarcalumenin is an under-investigated Ca^2+^-binding protein that is abundant within the SR and regulates Ca^2+^ reuptake by interacting with cardiac sarco(endo)plasmic reticulum Ca^2+^-ATPase2a (SERCA2a) (33). It has been reported to have a role in maintaining cardiac function during endurance exercise training and maintaining Ca^2+^ handling function of the SR in skeletal and cardiac cells and one of Sarcalumenin’s predicted partners, Atp2a1, has been proposed to interfere with hypoxia preconditioning and survival (38). Systemic ablation of *Sarcalumenin* caused enhanced resistance to muscle fatigue by compensatory changes in Ca^2+^ regulatory proteins that affect SOCE (store-operated Ca^2+^ entry) (39). Interestingly, human gene network analysis predicted interactions between *SRL* and *YWHAE*, which encodes 14-3-3 epsilon, the protein that sequesters cytoplasmic *CRTC* (10). In *C. elegans*, *col18* (*Sarcalumenin* ortholog) was shown to be 25x upregulated in glucose-fed animals (35) also suggesting a connection to CRTC signaling. In addition, our data suggests that abnormal calcium handling may play a key role in muscular dystrophy, where drastic reductions of Sarcalumenin in Dp427 (dystrophin of 427 kDa)-deficient fibers have been documented (40).

Our data in hiPSC-CMs, showing significant increases in the APD in response to KD of *CRTC* 2 and *CRTC* 3 (**Fig. 4**) as well as significant increases in SI in flies upon *tmn/SRL* KD and CRTC KD (**Fig. 8 A,B**) are all consistent with a Ca^2+^ overload hypothesis. Finally, concerted regulation of the Ca^2+^ binding protein Aralar (**Supplemental Table 5**) further suggests a role for CRTC in Ca^2+^ homeostasis. Taken together our data suggest that loss of *CRTC* and consequent reductions in *tmn / SRL* expression lead to a cytosolic Ca^2+^ overload (**Fig. 8D**) ultimately resulting in impaired contractility and mechanical and structural abnormalities in the heart. Taken together our data indicate that *CRTC* signaling plays a conserved and cardiac-autonomous role in maintaining heart structure and function. Thus, CaN - CRTC - Sarcalumenin may represent a parallel, perhaps conserved, pathway to that of CaN and NFAT in mediating cardiac hypertrophy and likely also play more general roles in muscle cell maintenance.

## MATERIALS AND METHODS

### Generation and maintenance of fly and Fish Stocks

*<u>Drosophila</u>* <u>stocks</u> were reared at 25°C on a standard laboratory diet consisting of yeast, corn starch, and molasses. All experiments were conducted on one or three-week old adult females. The *CRTC* mutant (*CRTC*) and UAS-*CRTC*(ii)-OE (overexpression) lines were a gift from Dr. Marc Montminy (Salk Institute). The *CREB* mutant (y*CREB*b[Δ400]/FM7c) was a gift from Dr. John Thomas (Salk Institute). Transgenic UAS-RNAi fly lines were obtained from the Vienna Drosophila Stock Center; the Bloomington Drosophila Stock Center; and the Kyoto Stock Center. The following Gal4 drivers were used: (cardiac) TinCΔ4-Gal4 (34) and Hand4.2-Gal4 (41); (pericardial cell) DOT-Gal4 (#BL6903) and SNS- Gal4 (#BL76160), (neurons) ELAV-Gal4 (#BL8765), (fat body) CG-Gal4 (#BL7011) and LSP2-Gal4 (#BL6357), and (muscle) MEF2-Gal4 (#BL50742, all BL are from Bloomington Stock Center, Bloomington, IN, USA). The following UAS-RNAi lines were used: *CRTC* (#V100974), Pp2B (#V103144), Calmodulin (#V102004), CRTC-Flag (#V318324), NFAT (#V107032, all #V from Vienna Stock Center, Biocenter 1, 1030 Wien, At.). The following UAS-OE lines were used: constitutively active Pp2B-14B (#DGRC116254, Drosophila Genomic Resource Center, 1001 E. 3rd St. Bloomington, IN). The following lines served as controls: w^1118^ (#BL3605), KK (#V100000), GD (#V35000), y[1]w[67c23] (#BL6599).

Homozygous *CREBb* mutant females were sterile and difficult to produce, thus, mutant males were also included in the experiments involving *CREBb* systemic mutants. (See **Supplemental Table 1** for a complete list of fly strains and crosses).

<u>Zebrafish stocks</u> (Oregon AB wild-type) were maintained under standard laboratory conditions at 28.5°C. Gene expression was manipulated using standard microinjection of morpholino (MO) antisense oligonucleotides (39). Subsequently, zebrafish were raised to 72 hours post fertilization (hpf); embryos were staged according to Kimmel et al. (32). All zebrafish experiments were performed in accordance with the protocols approved by SBP IACUC. For gene KD we used a previously characterized morpholino against CRTC3 (MO sequence: TCCTAATTTGGCTGAGCTTACCCTT, Gene Tools, LLC.) (38).

### *Drosophila* and zebrafish heart physiology

*Drosophila*: Electrical pacing assays were carried as previously described (17) . Intact adult flies were anesthetized with FlyNap (Carolina Biological Supply Co, Burlington, NC) placed between two rows of conductive gel overlying two electrodes and paced by applying a 40 V, 6 Hz square wave for 30 seconds. Abdominally located hearts were observed under a dissecting microscope; heart failure rate was defined as the percentage of hearts that did not beat or were visibly fibrillating 2 minutes after the end of the pacing regime.

Semi-intact preparations of the fly hearts were made as described previously (18) Briefly, flies were dissected in oxygenated artificial hemolymph (AHL) to expose the linear tube-like heart; excess fat was suctioned off with a micropipette. Following 15-20 minutes of recovery in fresh oxygenated AHL, 30- second high-speed movies (>140 fps) of contracting hearts were captured using via a Hamamatsu digital camera (EMCCD-C9300) on an Olympus BX61WI microscope with a 10x immersion objective. These movies were analyzed with the SOHA software (sohasoftware.com) (18, 19). The following key heart function parameters were measured: Diastolic Interval (DI), Systolic Interval (SI), Diastolic Diameter (DD), and Systolic Diameter (SD). Stroke Volume (SV) was estimated based on the assumption of a cylindrical heart tube using SV=(r^2^)*h, where h=1 and “r” is the tube radius and derived from the DD and SD measurements, i.e. SV =(1/2)DD)^2^ – (1/2SD)^2^.

Zebrafish: In-depth quantitative analyses of zebrafish cardiac function and conduction dynamics was performed as previously described (16). Larval zebrafish at 72 hpf were immobilized in a small volume of low melt agarose (1.5-2%) and submerged in conditioned water. Hearts were imaged in vivo at room temperature (20-21°C) with direct immersion optics and a digital high-speed camera (Orca Flash, Hamamatsu Photonics). High-speed movies (∼150 fps) were analyzed using SOHA )(33) to quantify heart period (R-R interval) and chamber size.

### Immunohistochemistry and optical sectioning

To assess morphological differences, immunohistochemistry was performed as described previously (42). Briefly, hearts were dissected, arrested/relaxed with 10mM EGTA and fixed for 20 min with 4% paraformaldehyde. The trimmed hearts were transferred to a 96-well plate (a maximum of ten hearts per well) and washed with PBSTx (phosphate-buffered saline, 0.1% Triton X-100). Dissected abdomens were incubated with primary antibodies diluted in PBTx for 2 hours at room temperature or overnight at 4°C and then washed in PBTx and incubated in secondary antibodies diluted in PBTx. Following 3 additional washes, hearts were mounted onto glass slides in Prolong Diamond anti-fade mounting medium with DAPI (ThermoFisher Scientific, Waltham, MA, USA). Z-stack images were obtained with a Zeiss ApoTome.2 and Zeiss Imager.Z1 Microscope system (Carl Zeiss, White Plains, NY, USA), at 10x and 25x magnification. Primary Antibodies used were: pericardin antibody (1:100, EC11, DSHB) to stain for the collagen-IV like extracellular matrix protein; α-actinin at 1:100 (gift from J. Saide), fly anti-CRTC antibody at 1:100 (M. Montminy), vertebrate CRTC antibody at 1:100 (MRC PPU Reagents and Services), anti-flag antibody (Sigma F3165), anti-H15/Nmr1 at 1:2000 (gift from J. Skeath (43), anti-Dystroglycan at 1:1000 (gift from A. Wodarz (44), Zfh1 at 1:1000 (45), anti-DMef2 at 1:2000 (gift from E. Olson (44), anti-Svp at 1:200 (gift from R. Cripps,(47). Secondary antibodies used were Alexa Fluor 488 donkey anti-sheep at 1:500 (Invitrogen, Carlsbad, CA, USA), Alexa Fluor 488 goat anti-mouse at 1:500 (Invitrogen, Carlsbad, CA, USA), Alexa Fluor 647 donkey anti-rabbit at 1:500 (Invitrogen, Carlsbad, CA, USA), Cy5 goat anti-guinea pig at 1:500 (Abcam, Cambridge, MA, USA). Alexa Fluor 594 phalloidin (Invitrogen, Carlsbad, CA, USA) was used to stain for F-actin or filamentous actin.

Optical sectioning for CM thickness measurements was done on both sides of the conical chamber ostia (anterior-most heart chamber). Optical sections were converted to binary images and the heart walls were identified using the Weka segmentation tool in ImageJ. Heart wall measurements were made at 4 points (2 dorsal and 2 ventral) for each of the 2 images per chamber and averaged. Care was taken to avoid measuring at locations containing nuclei, which expand the CM membranes.

Zebrafish hearts were fixed in 4% paraformaldehyde in PBS for 20 min and whole-mount immunofluorescence was performed as previously described (45, 46), using primary monoclonal antibodies against sarcomeric myosin heavy chain (MF20). MF20 was obtained from the Developmental Studies Hybridoma Bank maintained by the Department of Biological Sciences, University of Iowa. Secondary antibody, either donkey anti-chicken AlexaFluor488 (Jackson ImmunoResearch, 1:200) or Donkey anti-Rabbit AlexaFluor594 (Invitrogen, 1:200), was used in 1:200 dilution. Fluorescence images were acquired using an LSM 510 confocal microscope (Zeiss, Germany) with a 40X water objective.

### Climbing Assay

The negative geotaxis assays are modified from the RING (Rapid Iterative Negative Geotaxis) assays described in ((47)). Flies were sorted on CO_2_, by sex, into groups of twenty or less. After one hour of recovery time, they were transferred into polystyrene vials, marked with 1cm intervals. To induce the geotaxis response, each vial was then placed against a white background and tapped down firmly until all the flies fell to the bottom of the vial. Climbing ability/Negative geotaxis response was filmed with a digital camera over a ten second interval for each trial. Assays were performed three times for each set of flies with one minute recovery time in between each trial. For the analysis, video to jpeg conversion software was used to create still images at one second intervals for the 10 sec trial. The flies in each image were counted based on the 1 cm intervals marked on the vial. Flies that were found to be directly on a centimeter mark were counted in the higher distance bracket. Number of flies crossing the height of 2 and 10 cm were recorded. Measurements from all three trials were averaged.

### Triacylglyceride (TAG) Assay

Triacylglyerides were measured using the TAG assay as described previously )(48). Extracts obtained from three-week old intact female flies were used in a colorimetric assay whereby triacylglyerides are enzymatically cleaved into free fatty acids and glycerol, glycerol 3-phosphate and finally H_2_O_2_ which reacts with 4-aminoantipyrine (4-AAP) and 3,5-dichloro-2-hydroxybenzene sulfonate (3,5 DHBS) to produce a red colored product that was measured using a 96-well spectrophotometer. The level of TAGs in the samples was based on a standard curve generated using TAG standards (Stanbio Life Sciences) of known concentrations (0.0625 – 2 µg/µL). TAG content was normalized to total protein determined using a standard Bradford Protein assay (Bio-Rad Laboratories, Hercules, CA, USA).

### DNA extractions

DNA for PCR was extracted from whole flies using a standard organic extraction protocol. Materials included homogenizer tubes with polyacetal pestle (#K7496250030), Cell Lysis Solution (Qiagen #158906), RNase A Solution (Qiagen #158922), Protein Precipitation Solution (Qiagen #158910), DNA Hydration Solution (Qiagen #158914, QIAGEN, Germantown, MD, USA), 100% alcohol (Decon Laboratories, Inc.), and Isopropanol (ThermoFisher Scientific, Waltham, MA, USA). The final samples were assessed for quantity and quality using a NanoDrop spectrophotometer.

### RNA extractions

RNA was extracted from the hearts of either one-week old or three-week old female flies using the miRNeasy Mini Kit (QIAGEN, Germantown, MD, USA) as per manufacturer’s protocol. The final samples were assessed for quantity and quality using a NanoDrop spectrophotometer.

### RNA extraction, cDNA synthesis and quantitative PCR (Q-PCR)

Total RNA was isolated from fly hearts (15 per sample) in replicates of five with a RNeasy Isolation kit (QIAGEN, Germantown, MD, USA) per manufacturer’s protocol. Total RNA (25-30 ng) was quantified and checked for purity using a Nanodrop spectrophotometer. RNA was subsequently reverse-transcribed using a QuantiTect Reverse Transcription Kit (QIAGEN, Germantown, MD, USA), per manufacturer’s protocol. Perl Primer software (IDT) was used to design primers with optimal annealing temperatures and primer efficiency (**Supplemental Table 6**). The FastStart Essential DNA Green Master kit (Sigma-Aldrich Corp. St. Louis, MO, USA) was used to carry out qPCR in a Roche LightCycler® 96. RPL32 and GAPDH, and actin were used as reference genes.

Total RNA was extracted from zebrafish adult hearts using TRIzol (Invitrogen) and stabilized in RNA later (Thermo Fisher) and processed according to the RNeasy Micro Kit (Qiagen). Eight hearts were pooled together. RNA (1 µg) was reverse transcribed to cDNA with SuperScript reverse transcriptase (Invitrogen) using random hexamers. Q-PCR was performed as described previously(52). *β*-*actin* (*actb2*) and *GADPH* were used to normalize gene expression in the Q-PCR experiments. Primers for Q-PCR are listed in **Supplemental Table 6**. At least three independent biological replicates were performed.

### In situ hybridization

Gene expression in adult hearts was assessed for Gal4 using RNAscope (ACDbio, Newark, USA), and for CRTC and CG9297 using HCR (Molecular Instruments). Flies were dissected in oxygenated artificial hemolymph (AHL) to expose the linear tube-like heart; excess fat was suctioned off with a micropipette, and hearts were fixed for 20 minutes in 4% paraformaldehyde. Hybridization and fluorescent labeling was performed according to manufacturers’ protocols.

### RNA Sequencing and Analysis

Hearts from 15 female flies were dissected as described previously, removed with fine forceps and pooled as a single sample. RNA was extracted using the Qiagen RNAeasy mini kit (QIAGEN, Germantown, MD, USA). Six replicate samples were obtained from 1-week old wildtype (w^1118^) and *CRTC*^-/-^ knockout flies. Six to seven replicate samples were obtained from 3-week old TincΔ4-Gal4>UAS-*CRTC*-RNAi, TincΔ4-Gal4>UAS-*CRTC*-OE, and TincΔ4-Gal4>KK-control flies. PolyA RNA was isolated using the NEBNext® Poly(A) mRNA Magnetic Isolation Module and barcoded libraries were made using the NEBNext® Ultra II™ Directional RNA Library Prep Kit for Illumina® (NEB, Ipswich, MA, USA).

Libraries were pooled and paired-end sequenced (2X75) on the Illumina NextSeq 500 using the High output V2.5 kit (Illumina Inc., San Diego, CA, USA). Average read count for w^1118^ and *CRTC* mutant transcripts was approximately 15.3 million reads per sample, as 26 samples were shared on a single flow cell with maximum read count of 400 million reads. Average read count for *CRTC*-RNAi, *CRTC*-OE, and Control^KK^ was 19 million reads per sample, as 21 samples were shared on a single flow cell with maximum read count of 400 million reads. Read data were processed in BaseSpace (basespace.illumina.com). Reads were aligned to *Drosophila melanogaster* genome (Dm6) using STAR aligner (https://code.google.com/p/rna-star/) with default settings. Differential transcript expression was determined using the Cufflinks Cuffdiff package (https://github.com/cole-trapnell-lab/cufflinks).

### iPSC-CM Assay

#### Generation of hiPSC-CMs

Id1 overexpressing hiPSCs (45, 49) were dissociated with 0.5 mM EDTA (ThermoFisher Scientific, Waltham, MA, USA) in PBS without CaCl2 and MgCl2 (Corning) for 7 min at room temperature. hiPSC were resuspended in mTeSR-1 media (StemCell Technologies, Cambridge, MA 02142) supplemented with 2 µM Thiazovivin (StemCell Technologies, Cambridge, MA 02142) and plated in a Matrigel-coated 12-well plate at a density of 3 x 105 cells per well. After 24 hours after passage, cells were fed daily with mTeSR-1 media (without Thiazovivin) for 3-5 days until they reached ≥ 90% confluence to begin differentiation. hiPSC-CMs were differentiated as previously described (40, 41). At day 0, cells were treated with 6 µM CHIR99021 (Selleck Chemicals, Houston, TX, USA) in S12 media (42) for 48 hours. At day 2, cells were treated with 2 µM Wnt-C59 (Selleck Chemicals, Houston, TX, USA), an inhibitor of WNT pathway, in S12 media. 48 hours later (at day 4), S12 media is fully changed. At day 5, cells were dissociated with TrypLE Express (ThermoFisher Scientific, Waltham, MA, USA) for 2 min and blocked with RPMI supplemented with 10% FBS (Omega Scientific, Tarzana, CA, USA). Cells were resuspended in S12 media supplemented with 4 mg/L Recombinant Human Insulin (ThermoFisher Scientific, Waltham, MA, USA) (S12+ media) and 2 µM Thiazovivin and plated onto a Matrigel-coated 12-well plate at a density of 9 x 105 cells per well. S12+ media was changed at day 8 and replaced at day 10 with RPMI (ThermoFisher Scientific, Waltham, MA, USA) media supplemented with 213 µg/µL L-ascorbic acid (Sigma-Aldrich, St. Louis, MO, USA), 500 mg/L BSA-FV (Gibco), 0.5 mM L-carnitine (Sigma-Aldrich, St. Louis, MO, USA) and 8 g/L AlbuMAX Lipid-Rich BSA (CM media, ThermoFisher Scientific, Waltham, MA, USA). Typically, hiPSC-CMs start to beat around day 9-10. At day 15, cells were purified with lactate media (RPMI without glucose, 213 µg/µL L-ascorbic acid, 500 mg/L BSA-FV and 8 mM Sodium-DL-Lactate (Sigma-Aldrich, St. Louis, MO, USA), for 4 days. At day 19, media was replaced with CM media.

### Voltage assay in hiPSC-CMs

Voltage assay is performed using labeling protocol described in (50). Briefly, hiPSC-CMs at day 25 of differentiation were dissociated with TrypLE Select 10X for up to 10 min and action of TrypLE was neutralized with RPMI supplemented with 10% FBS. Cells were resuspended in RPMI with 2% KOSR (Gibco) and 2% B27 50X with vitamin A (Life Technologies, Carlsbad, CA, USA) supplemented with 2 µM Thiazovivin and plated at a density of 6,000 cells per well in a Matrigel-coated 384-well plate. hiPSC-CMs were then transfected with siRNAs directed against each *CRTC* complex components (ON-TARGETplus SMART pool, si*CRTC*1: L-014026-01-0005, si*CRTC*2: L- 018947-00-0005, si*CRTC*3: L-014210-01-0005) using lipofectamine RNAi Max (ThermoFisher Scientific, Waltham, MA, USA). Each siRNA was tested individually and in combination in 8-plicates. Three days post-transfection, cells were first washed with pre-warmed Tyrode’s solution (Sigma-Aldrich, St. Louis, MO, USA) by removing 50 µL of media and adding 50 µL of Tyrode’s solution 5 times using a 16-channel pipette. After the fifth wash, 50 µL of 2x dye solution consisting in voltage sensitive dye Vf2.1 Cl (Fluovolt, 1:2000, ThermoFisher Scientific, Waltham, MA, USA) diluted in Tyrode’s solution supplemented with 1 µL of 10% Pluronic F127 (diluted in water, ThermoFisher Scientific, Waltham, MA, USA) and 20 µg/mL Hoescht 33258 (diluted in water, ThermoFisher Scientific, Waltham, MA, USA) was added to each well. The plate was placed back in the 37°C 5% CO2 incubator for 45 min. After incubation time, cells were washed 4 times with fresh pre-warmed Tyrode’s solution using the same method described above. hiPSC-CMs were then automatically imaged with ImageXpress Micro XLS microscope at an acquisition frequency of 100 Hz for a duration of 5 sec with excitation wavelength of 485/20 nm and emission filter 525/30 nm. A single image of Hoescht was acquired before the time series. Fluorescence over time quantification and trace analysis were automatically quantified using custom software packages developed by Molecular Devices and Colas lab. Two independent experiments were performed. KD efficiency was confirmed 2 days post-siRNA-transfection by qPCR (see **Supplemental Table 6** for primer sequences).

### RNA extraction and qPCR Assay with hiPSC-CMs

The hiPSC-CMs were dissociated at day 25 of differentiation with TrypLE Select 10X for up to 10 min and action of TrypLE was neutralized with RPMI supplemented with 10% FBS. Cells were resuspended in RPMI (ThermoFisher Scientific, Waltham, MA, USA) media supplemented with 213 µg/µL L-ascorbic acid (Sigma-Aldrich, St. Louis, MO, USA), 500 mg/L BSA-FV (Gibco), 0.5 mM L-carnitine (Sigma-Aldrich, St. Louis, MO, USA) and 8 g/L AlbuMAX Lipid-Rich BSA (CM media, ThermoFisher Scientific, Waltham, MA, USA) supplemented with 2 µM Thiazovivin and plated at a density of 6,000 cells per well in a Matrigel-coated 384-well plate. hiPSC-CMs were then transfected with siRNAs directed against each *CRTC* complex components (ON-TARGETplus SMART pool, si*CRTC*1: L-014026-01-0005, si*CRTC*2: L-018947-00-0005, si*CRTC*3: L-014210-01-0005) using lipofectamine RNAi Max (ThermoFisher Scientific, Waltham, MA, USA). Each siRNA was tested individually and in combination in 4-plicates. Three days post-transfection, total RNA was extracted using Zymo Research Quick-RNA MircoPrep Kit (Zymo Research, R1051) following the manufacturers’ recommendations. RNA concentration was measured by Nanodrop (Thermo Scientific). Aliquots of 1 μg of RNA were reverse transcribed using a QuantiTect Reverse Transcription kit (Qiagen, 205314), and qPCR was performed with iTaq SYBR Green (Life Technologies) using a 7900HT Fast Real-Time PCR system (Applied Biosystems). Gene expression was normalized to that of glyceraldehyde 3-phosphate dehydrogenase (GAPDH) for human iPSC-dervied cardiomyocyte samples using the 2−ΔΔCt method. Human primer sequences for qRT-PCR were obtained from Harvard Primer Bank (see **Supplemental Table 6**).

### *In silico* TFBS analysis in R

Binding site motifs for *CrebB-17A* and *CrebA* were imported from MotifDB (51) and searched for in aligned genome assemblies of *D. melanogaster/persimilis* and *D.melanogaster/yakuba*. This identified all sites that were conserved between those genome pairs. Next, this was further narrowed down to those sites that localize within a -2000bp/+1000bp window of annotated transcriptional start sites (*dm6* assembly). The R script for this analysis is available at https://github.com/gvogler/Dondi_2023.

### Statistical Analysis

Fly and fish cardiac function data was analyzed using the D’Agostino and Pearson omnibus normality test for Gaussian distribution. For normally distributed data, statistical significance was determined using a 1-way ANOVA for simple comparisons between more than two groups and 2-way ANOVA (for multiple manipulations) followed by multiple comparisons post-hoc tests as indicated in Figure legends. Data sets that did not show a normal distribution were analyzed using a nonparametric 2-tailed unpaired t-test, Wilcoxcon Rank Sum test, or Kruskal-Wallis test followed by Dunn multiple comparisons post-hoc tests.

For hiPSC-ACM data we used the nonparametric Kolmogorov-Smirnov test to compare the differences in the cumulative distributions of APD data. Population distribution of control and siRNA-treated hiPSC-CMs was generated with GraphPad Prism software (2019) using nonlinear regression. To determine statistical significance between experimental and control groups, we used two-tailed, unpaired Student’s t-test. All statistical analyses were performed using GraphPad Prism software.

## Acknowledgements

We thank Ms. Erika Taylor, Mr. Arthur Bautista, Mr. Sean Paknoosh, and Dr. Xin-Xin I. Zeng for technical assistance.

## Sources of Funding

This work was supported by:

RB - NIH R01 HL149992 and NIH R01 HL054732 AC - R01 HL153645 and R01 HL148827

AG - NIH grant 5F31HL134305-02

MM - NIH grant R01 DK083834, the Leona M. and Harry B. Helmsley Charitable Trust, the Clayton Foundation for Medical Research, and the Kieckhefer Foundation.

KO - The Sanford Burnham Prebys Medical Discovery Institute, the American Heart Association (14GRNT20490239).

JBT - NIH grant R01DK077979.

---

**Supplemental Figure 1.**
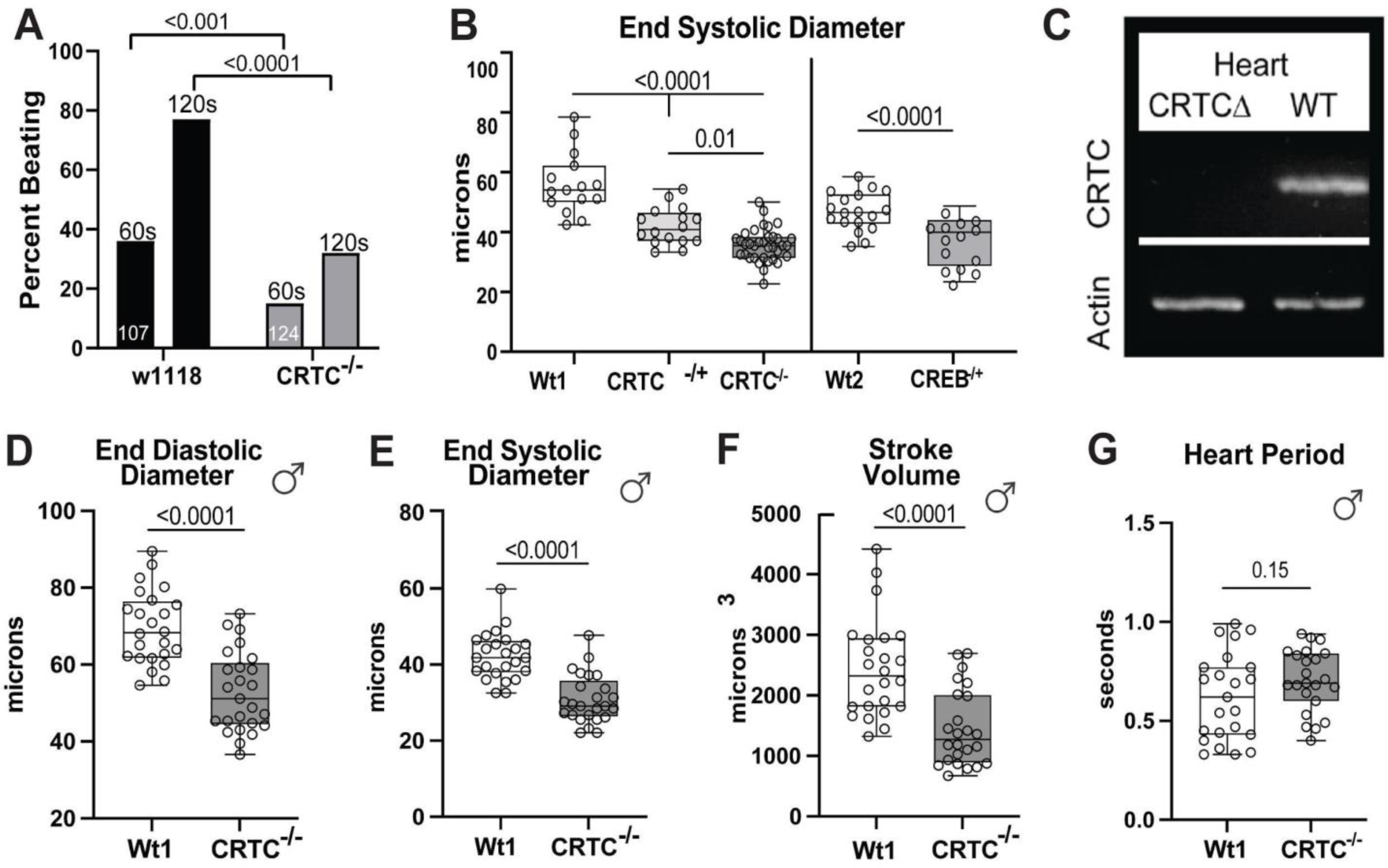
Deletion of *CRTC* makes flies more susceptible to stress-induced heart failure. **(A)** Heart Failure assay showing the percent of hearts still beating following a 30 sec electrical pacing stress test. Approximately 60% of hearts in wt flies (w1118, N=107) were not beating 60s after electrical pacing (6HZ for 30s); all but ∼20% resumed beating by 120s post stress. Roughly 80% of hearts in *CRTC* mutants (N=124) were not beating 60s post pacing stress and more than 60% were still not beating at 120s. Significance determined by Multiple Mann-Whitney tests. **(B)** End Systolic diameters of hearts from 1 week old flies. Hearts from both *CRTC* (left) and *CREB* mutants (right) had significantly reduced diameters compared to Wt controls. Significance was determined by 1-way ANOVA for Wt1 and *CRTC* mutants with Sidaks multiple comparison post hoc test. and unpaired student t-test for Wt2 and *CREB* mutant. (Max, Min and median with p values are shown.) **(C)** qPCR of isolated hearts show robust expression in wildtype (WT) controls but not in hearts from *CRTC* mutants. **(D)** End diastolic diameters, **(E)** end systolic diameters, and **(F)** stroke volume were all reduced in hearts from 1 week old *CRTC* mutant males compared to Wt controls. **(G)** Heart period was unaffected by *CRTC* KO in males (D-G, unpaired student t-test, plots show Max, Min and Median, p values shown).

**Supplemental Figure 2.**
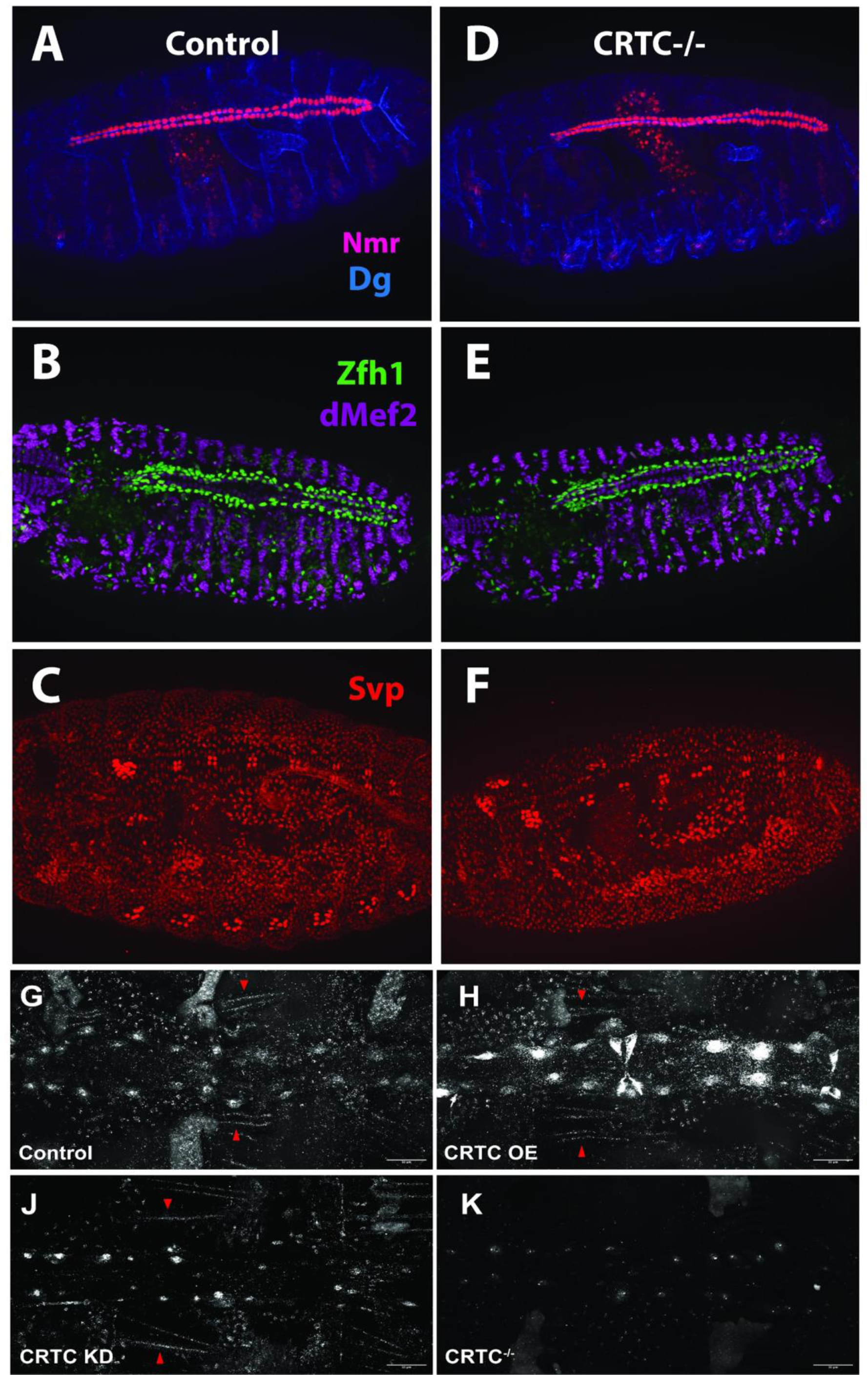
Systemic *CRTC* KO did not affect cardiac development. **(A-C)** Stage 17 wt and **(D-F)** *CRTC* mutant embryos. were examined for cardiac development. (**A,D)** Wildtype and CRTC mutant embryos were stained for Neuromancer (Nmr) which labeled cardioblast nuclei (red), and dystroglycan (Dg) which labels the basal domain of epithelial cells (blue). ( **B,E)** Zfh1 (green) labeled the nuclei of pericardial cells and dMEF2 (purple) stained all muscle nuclei. ( **C,F)** Svp labels a subset of cardiac cells that will form the ostia (inflow tract). **(G)** HCR for CRTC shows CRTC mRNA expression in cardiac tube, nuclei and somatic muscle in control fly **(H)** In cardiac specific OE of CRTC (TinCd4-Gal4 driver), HCR confirms the increase of CRTC mRNA in the cardiac tube and cardiomyocytes nuclei. CRTC mRNA expression stays the same in somatic muscles (arrowheads) **(J)** HCR staining shows the decreased CRTC mRNA in the cardiac tube and cardiomyocytes nuclei in response to cardiac specific CRTC KD (TinCd4-Gal4 driver). CRTC mRNA expression stays the same in somatic muscles (arrowheads). **(K)** In CRTC-/- flies, HCR shows the dramatic loss of CRTC mRNA in every tissue.

**Supplemental Figure 3.**
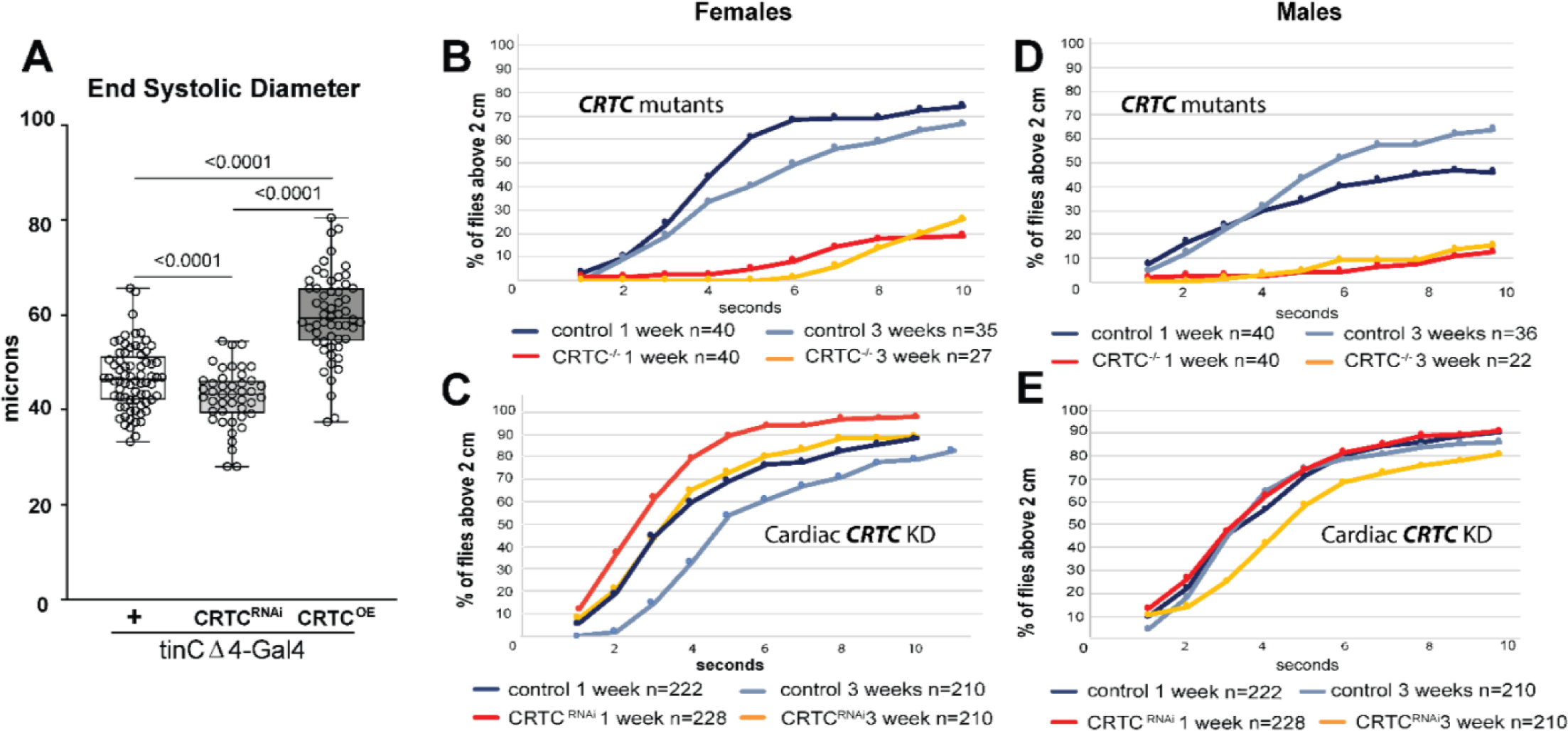
CRTC affects Somatic muscle function. **(A)** End Systolic Diameters in response to cardiac-specific CRTC KD. (Graph shows Max, Min and Median values; significance determined by one-way ANOVA, p values are shown.) **(B - E)** Climbing assay results; all flies assayed at 1 or 3 weeks. **(B)** Female control flies (blue traces) and *CRTC systemic mutants* (orange traces) were assayed and the percent of flies above the 2 cm mark in the assay vial over a 10 sec interval is shown. **(C)** Percent of female control flies (blue traces) and *cardiac-specific CRTC KD* flies (orange traces) above the 2 cm mark over a 10 sec interval is shown. **(D)** Percent of male control flies (blue traces) and *CRTC systemic mutant flies* (orange traces) the 2 cm mark in the assay vial over a 10 sec interval is shown. **(E)** Percent of male control flies (blue traces) and *cardiac-specific CRTC KD* flies (orange traces) above the 2 cm mark over a 10 sec interval is shown.

**Supplemental Figure 4.**
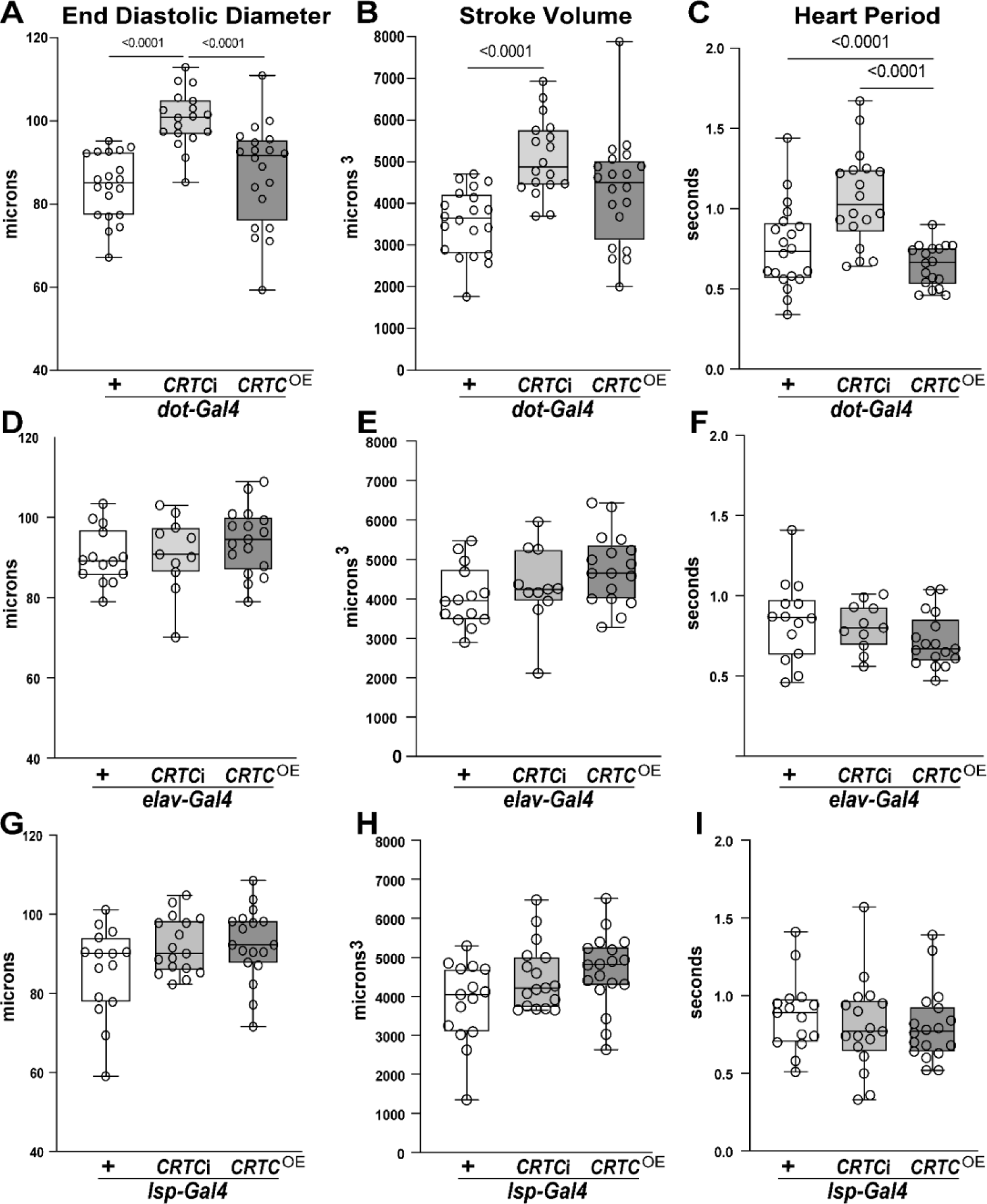
*CRTC* cardiac phenotype is not recapitulated by *CRTC* KD in nephrocytes, neurons, or fat body. **(A)** End Diastolic Diameter (EDD) and **(B)** stroke volume were increased by nephrocyte-specific KD of *CRTC*, using the Dot-Gal4 driver. **(C)** Heart period was decreased (increased rate) by nephrocyte-specific *CRTC* OE. **(D)** EDD and **(E)** stroke volume and **(F)** heart period were unaffected by neuronal KD or OE of *CRTC*, using the elav-Gal4 driver. **(G)** EDD, **(H)** stroke volume and **(I)** heart period were unaffected by fat body-specific *CRTC* KD or OE. Plots show Max, Min and Median, significance was determined using a one-way ANOVA with Tukey’s multiple comparisons post hoc test, p values shown.

**Supplemental Figure 5.**
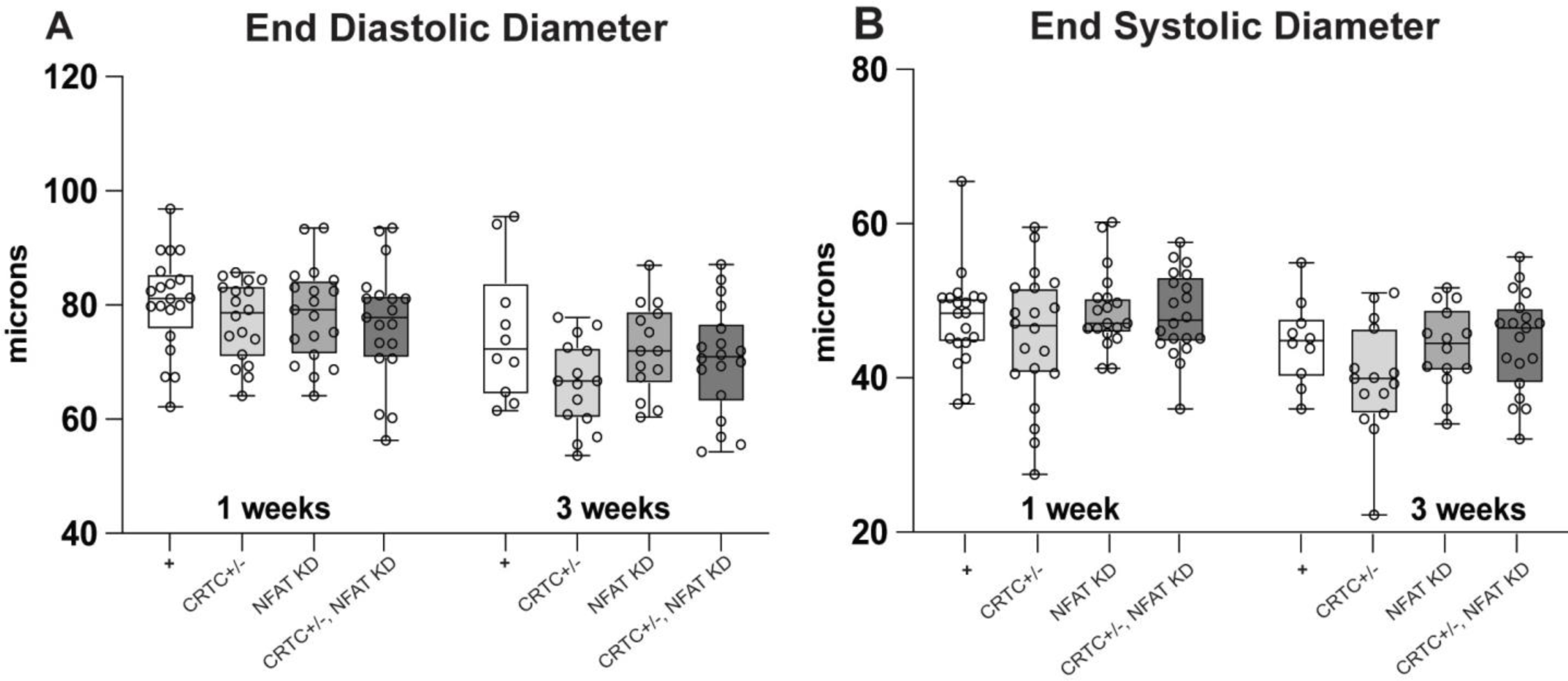
Cardiac KD of *NFAT* had little effect on heart function. **(A)** End Diastolic Diameters (ESD) were unchanged in response to cardiac NFAT KD with tinCΔ4-Gal4. **(B)** End Systolic Diameters (EDD) in hearts from flies with cardiac *NFAT* KD were unchanged. Loss of function of *CRTC* and *NFAT* together doesn’t show significant effect suggesting the absence of interaction between these two factors in the heart. Plots show Max, Min and Median, significance was determined using a one-way ANOVA with Tukey’s multiple comparisons post hoc test, p values shown.

**Supplemental Figure 6.**
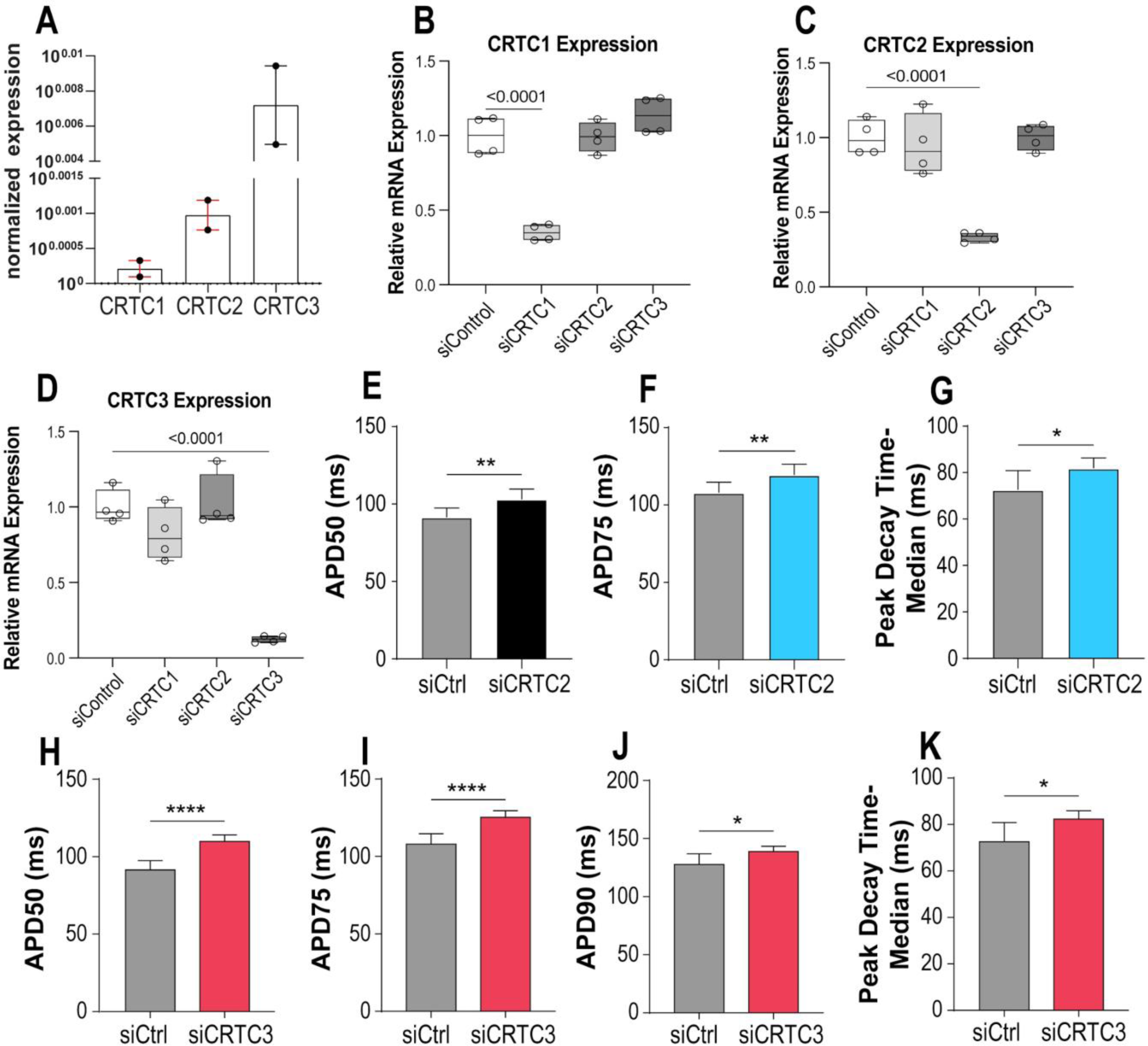
*CRTC 2 & 3* affect cardiac function in zebrafish and hiPSC-Cardiomyocytes (hiPSC-CM). **(A)** *CRTC* expression in isolated adult (10 month old) zebrafish hearts, normalized to elongation factor eF1a, showed significant expression of *CRTC 3* compared to *CRTC 1 & 2*. Data points represent 2 biological samples, 8 hearts per sample done in triplicate. **(B-D)** Relative mRNA expression shows that siRNA against CRTC1,2&3 efficiently KD the specific isoform in hIPSC-CMs **(E-K)** siRNA KD of *CRTC 2 &* 3 in hiPSC-CMs caused APD broadening and delayed repolarization Action Potential Duration was significantly increased with siRNA-mediated KD of *CRTC 3* at **(H)** APD50 **(I)** APD75 and **(J)** APD90 (i.e. at 50%, 75% and 90% repolarization, respectively). **(E)** APD50 and **(F)** APD75 were significantly broadened in response to siRNA-mediated KD of *CRTC2*. **(K)** *CRTC3* KD and **(G)** *CRTC2* both caused significant increases in peak decay times. Significance determined by unpaired t-test, *p<0.05, **p<0.01, ****P<0.0001.

**Supplemental Figure 7.**
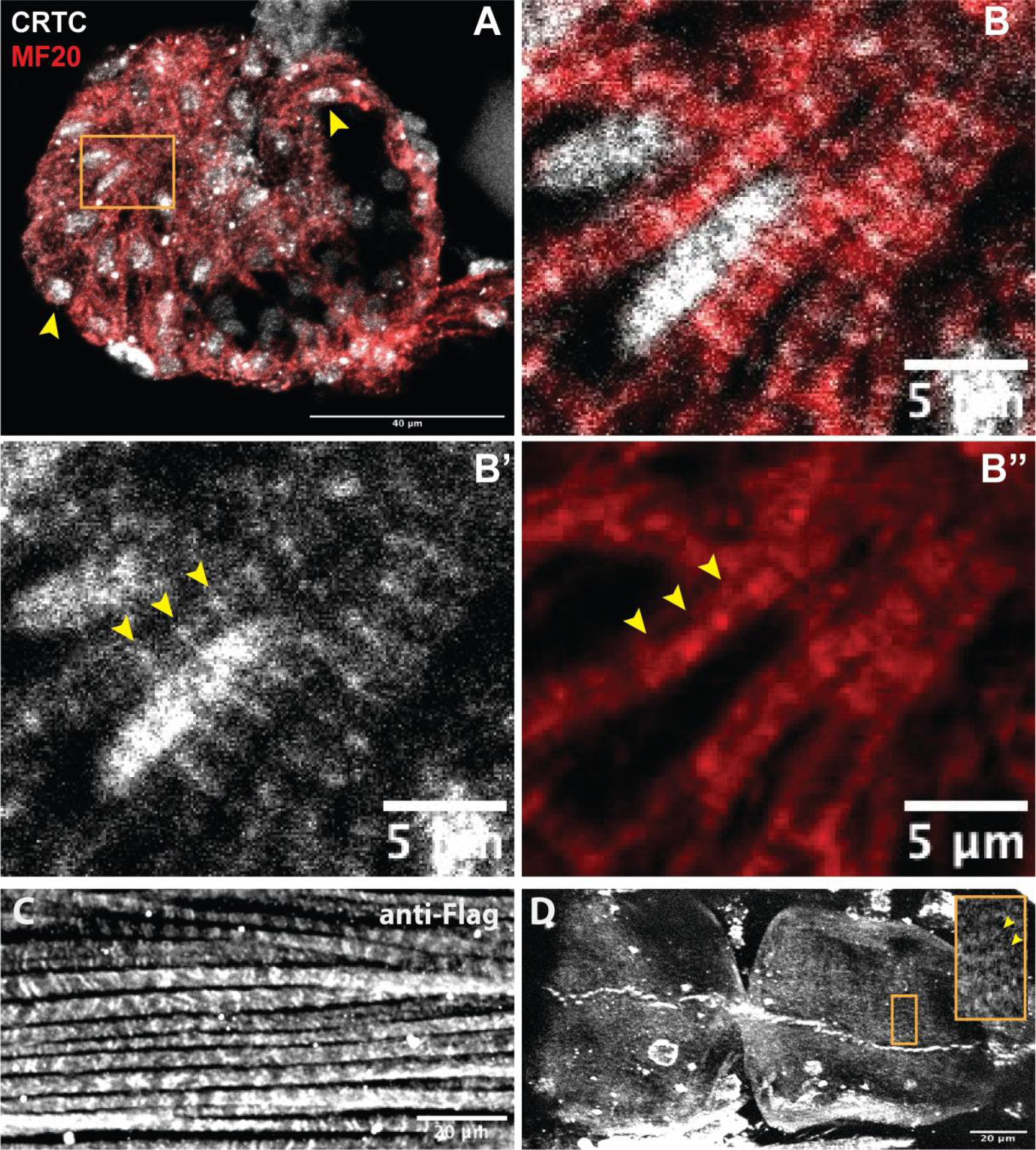
CRTC localizes to Z bands in zebrafish hearts. **(A)** CRTC (white) and myosin heavy chain (MY20, red) staining in zebrafish hearts. Arrowheads show CRTC localization in the nuclei. Scale bar is 40 µm. **(B, B’, B”’)** Higher magnification of the region in the yellow rectangle. (B’) Arrows show CRTC localization in a banded pattern outside the nucleus. (B”) Heavy MHC staining on either side of the CRTC bands suggests CRTC localizes to Z-bands. Scale bar is 5 µm.**(C)** Anti-Flag staining (white) of hearts from a CRTC Flag-tagged transgenic line confirmed CRTC localization in non-myocardial, ventral longitudinal fibers. **(D)** Optical section through a chamber of the fly heart tube stained with anti-Flag. On the right, higher magnification of the region in the yellow. Arrowheads show CRTC localization with anti-Flag staining around the Z-bands region in the sarcomere. Scale bars are 20 µm.

**Supplemental Figure 8.**
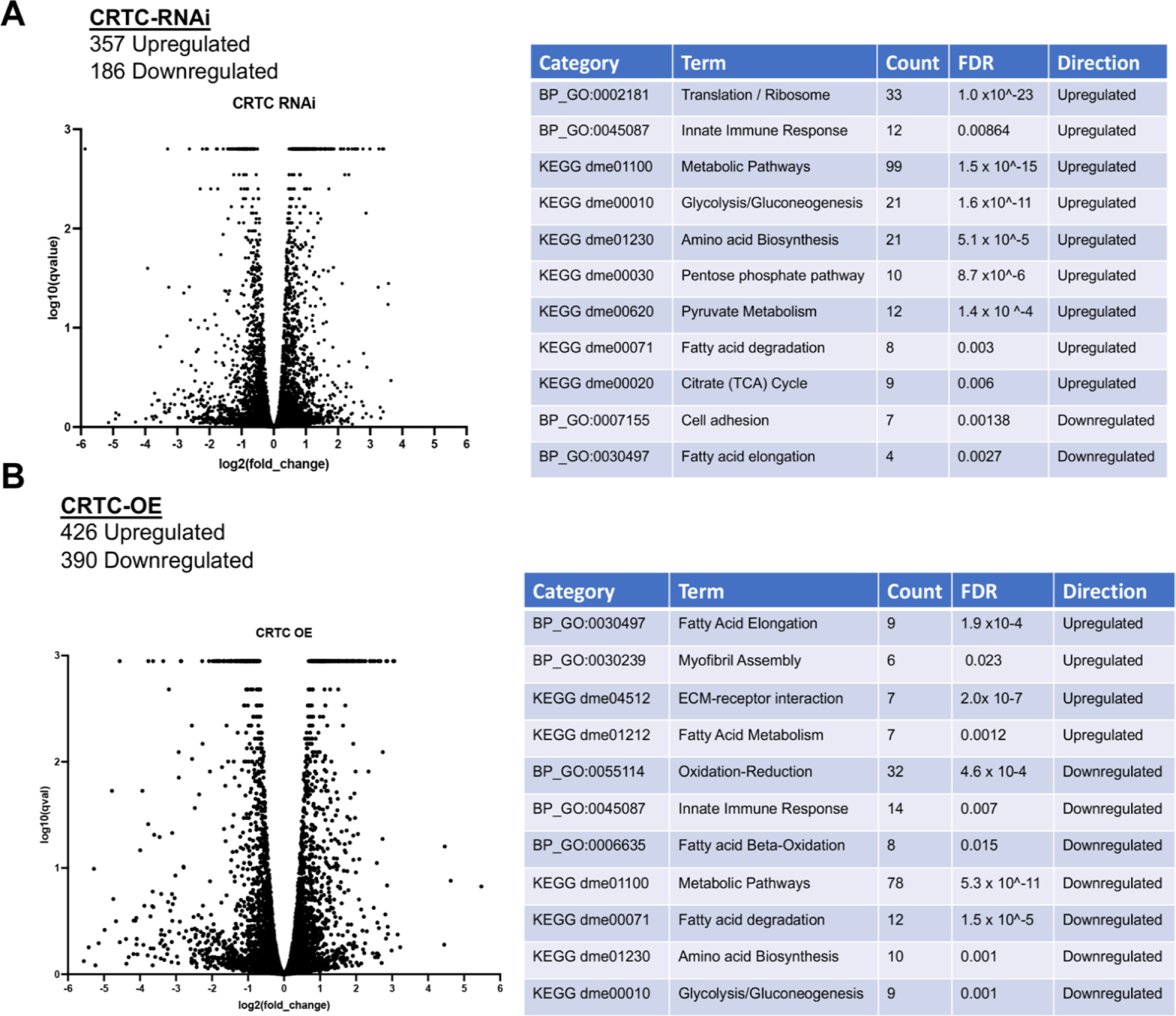
Cardiac *CRTC* KD and OE concertedly regulate metabolism in the heart. **(A, left)** Volcano plot of genes differentially regulated in hearts with cardiac-specific *CRTC* KD. **(A, right)** Differentially regulated genes fall into GO categories primarily related to cell metabolism. **(B, left)** Volcano plot of genes differentially regulated in hearts with cardiac-specific *CRTC* OE. **(B, right)** Differentially regulated genes fall into GO categories primarily related to cell metabolism.

**Table 1.**
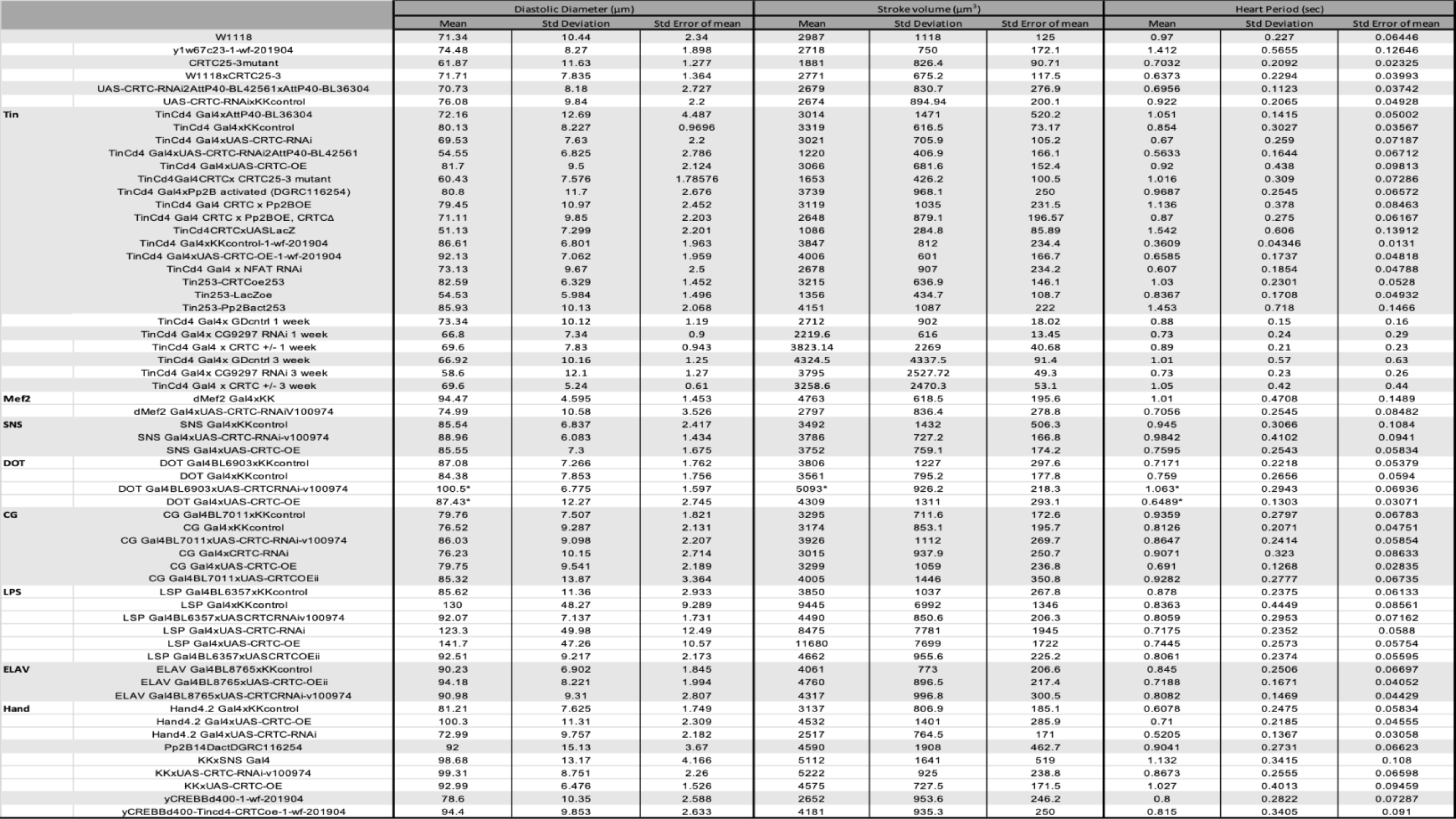
Complete list of fly crosses and heart function data.

**Table 2.**
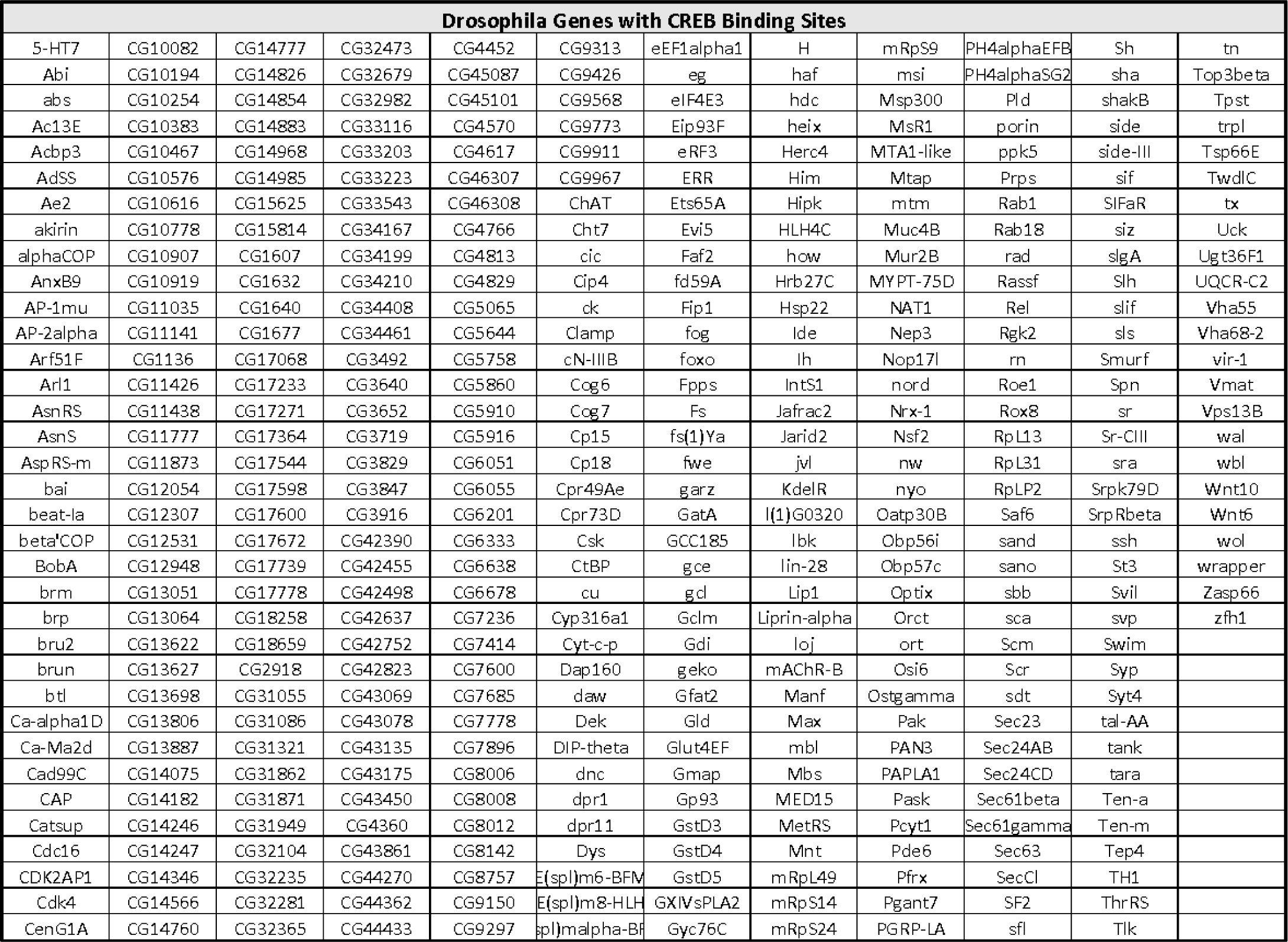
Fly genes with CREB binding sites conserved between *Drosophila melanogaster* and *Drosophila persimilis*.

**Table 3.** Hypergeometric test for CRTC/CREB-enriched target genes.

|  | Differential<br>expression | Not differentially<br>expressed | total |
| --- | --- | --- | --- |
| with CREB-TFBS | <u>101</u> | 814 | 915 |
| No CREB-TFBS | 1074 | 11938 | 13912 |
| total | 1175 | 12752 | 13927 |

**Contingency table for genes with CREB TFBS and differential expression after CRTC KD -or- OE.** Hypergeometric test for enrichment analysis using the values from the contingency table indicates that genes with CREB TFBS are more likely found to be overexpressed than genes without.

**Table 4.**
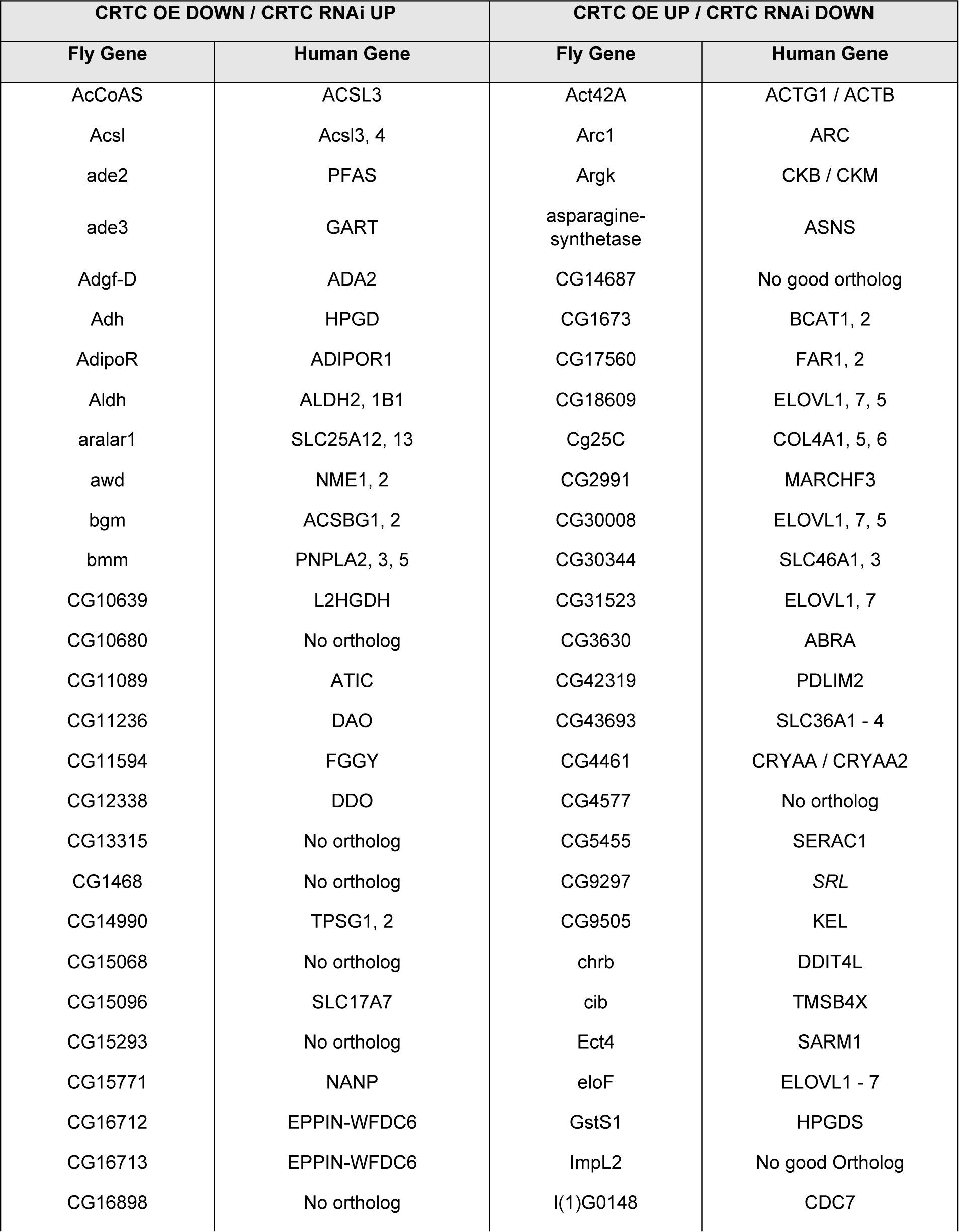
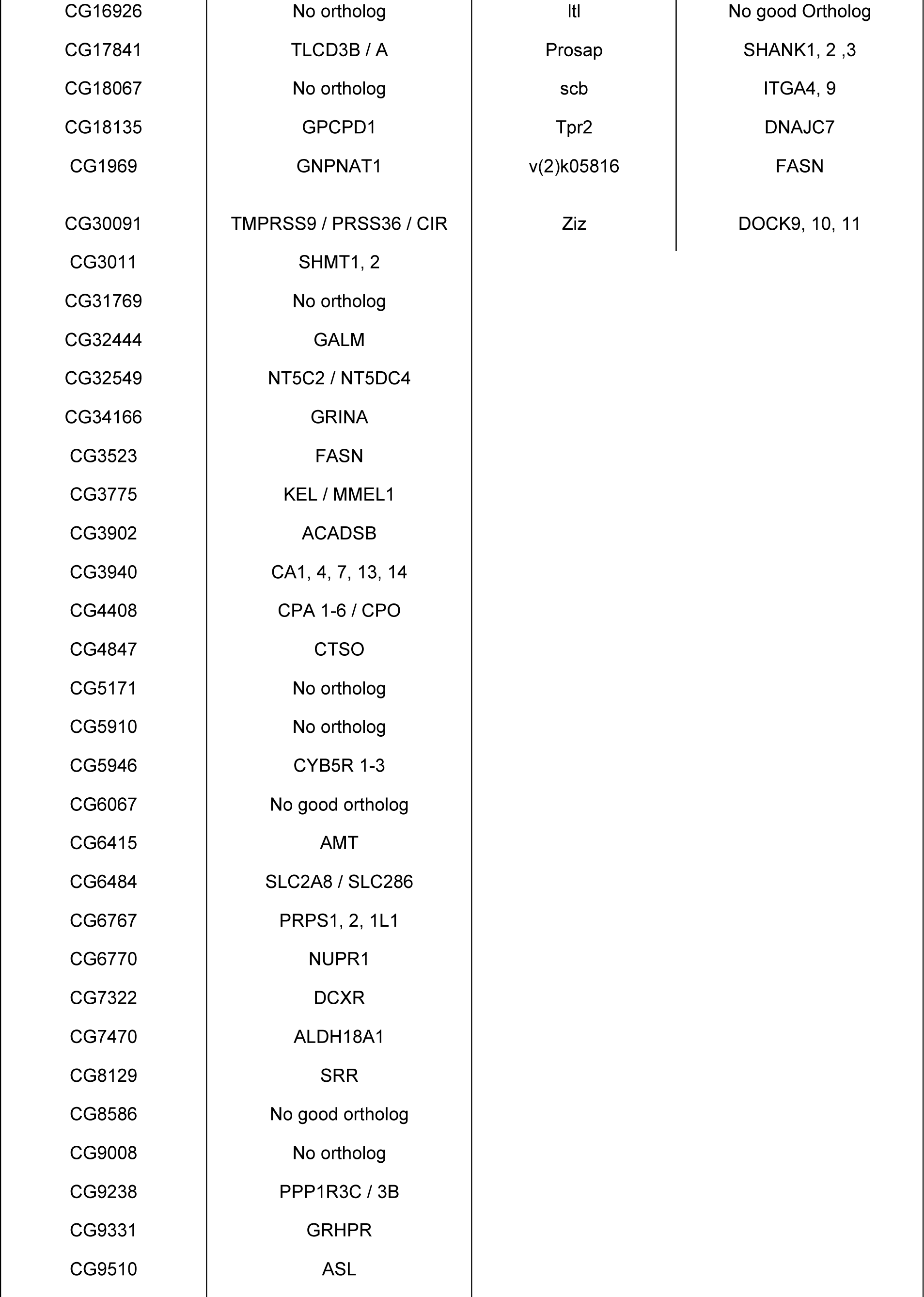
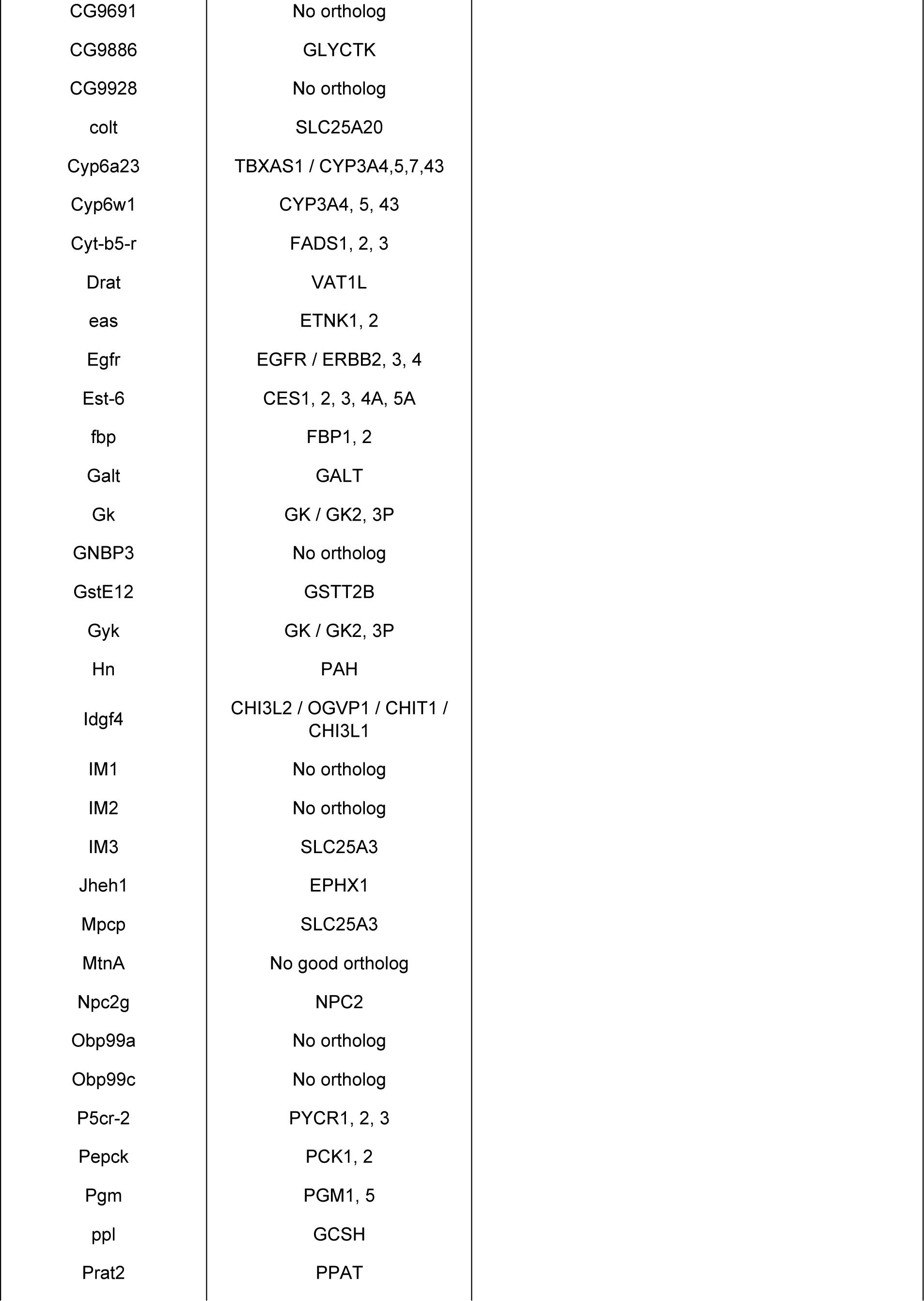
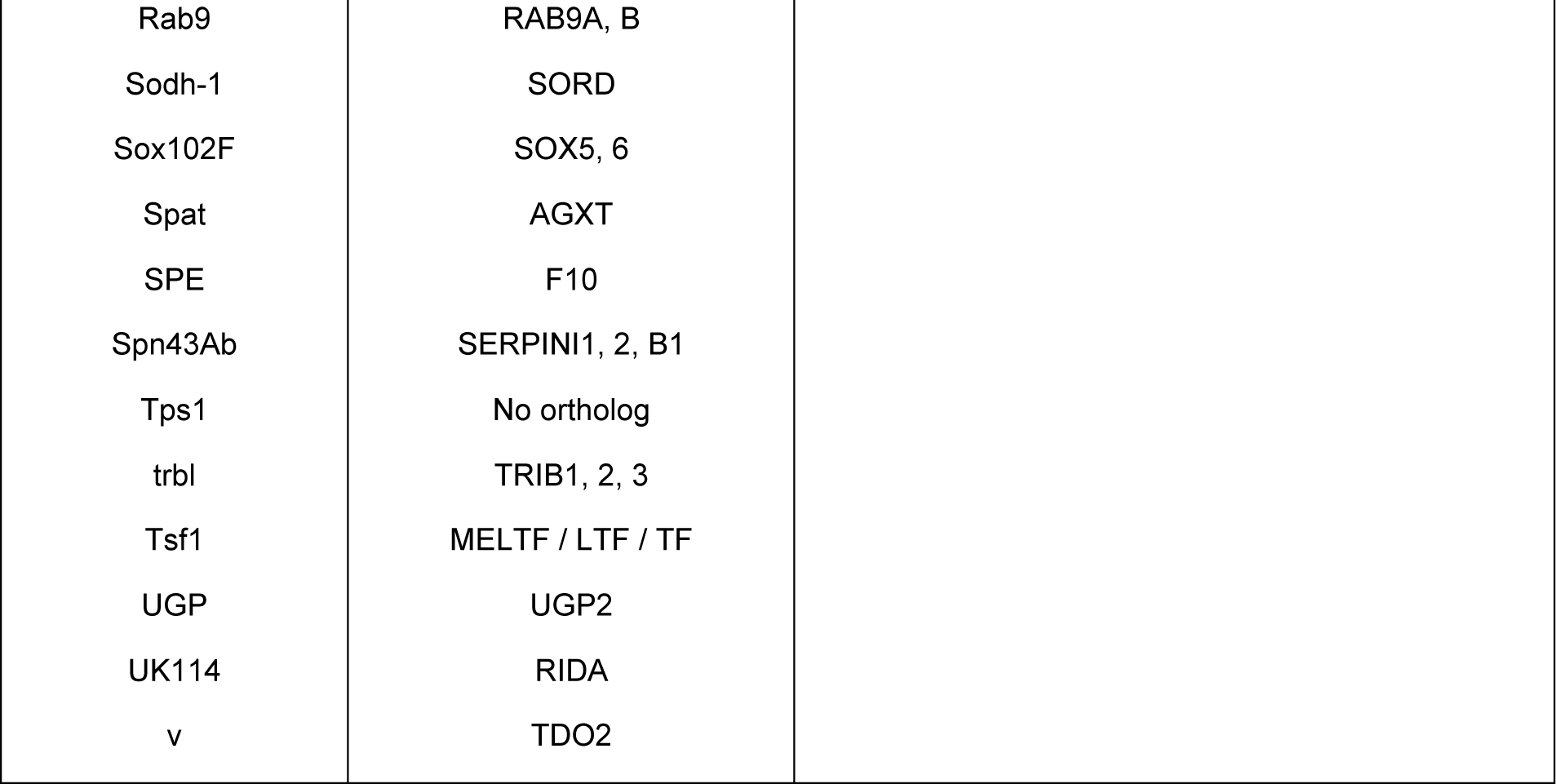
Complete list of Contra-regulated genes and human orthologs. Orthologous genes were identified using DIOPT - DRSC Integrative Ortholog Prediction Tool (v. 9). (https://www.flyrnai.org/cgi-bin/DRSC_orthologs.pl)

| CRTC OE DOWN / CRTC RNAi UP |  | CRTC OE UP / CRTC RNAi DOWN |  |
| --- | --- | --- | --- |
| Fly Gene | Human Gene | Fly Gene | Human Gene |
| AcCoAS | ACSL3 | Act42A | ACTG1 / ACTB |
| AcsI | AcsI3, 4 | Arc1 | ARC |
| ade2 | PFAS | Argk | CKB / CKM |
| ade3 | GART | asparagine-synthetase | ASNS |
| Adgf-D | ADA2 | CG14687 | No good ortholog |
| Adh | HPGD | CG1673 | BCAT1, 2 |
| AdipoR | ADIPOR1 | CG17560 | FAR1, 2 |
| Aldh | ALDH2, 1B1 | CG18609 | ELOVL1, 7, 5 |
| aralar1 | SLC25A12, 13 | Cg25C | COL4A1, 5, 6 |
| awd | NME1, 2 | CG2991 | MARCHF3 |
| bgm | ACSBG1, 2 | CG30008 | ELOVL1, 7, 5 |
| bmm | PNPLA2, 3, 5 | CG30344 | SLC46A1, 3 |
| CG10639 | L2HGDH | CG31523 | ELOVL1, 7 |
| CG10680 | No ortholog | CG3630 | ABRA |
| CG11089 | ATIC | CG42319 | PDLIM2 |
| CG11236 | DAO | CG43693 | SLC36A1 - 4 |
| CG11594 | FGGY | CG4461 | CRYAA / CRYAA2 |
| CG12338 | DDO | CG4577 | No ortholog |
| CG13315 | No ortholog | CG5455 | SERAC1 |
| CG1468 | No ortholog | CG9297 | SRL |
| CG14990 | TPSG1, 2 | CG9505 | KEL |
| CG15068 | No ortholog | chrb | DDIT4L |
| CG15096 | SLC17A7 | cib | TMSB4X |
| CG15293 | No ortholog | Ect4 | SARM1 |
| CG15771 | NANP | eloF | ELOVL1 - 7 |
| CG16712 | EPPIN-WFDC6 | GstS1 | HPGDS |
| CG16713 | EPPIN-WFDC6 | ImpL2 | No good Ortholog |
| CG16898 | No ortholog | I(1)G0148 | CDC7 |
| CG16926 | No ortholog | Itl | No good Ortholog |
| CG17841 | TLCD3B / A | Prosap | SHANK1, 2 ,3 |
| CG18067 | No ortholog | scb | ITGA4, 9 |
| CG18135 | GPCPD1 | Tpr2 | DNAJC7 |
| CG1969 | GNPNAT1 | v(2)k05816 | FASN |
| CG30091 | TMPRSS9 / PRSS36 / CIR | Ziz | DOCK9, 10, 11 |
| CG3011 | SHMT1, 2 |  |  |
| CG31769 | No ortholog |  |  |
| CG32444 | GALM |  |  |
| CG32549 | NT5C2 / NT5DC4 |  |  |
| CG34166 | GRINA |  |  |
| CG3523 | FASN |  |  |
| CG3775 | KEL / MMEL1 |  |  |
| CG3902 | ACADSB |  |  |
| CG3940 | CA1, 4, 7, 13, 14 |  |  |
| CG4408 | CPA 1-6 / CPO |  |  |
| CG4847 | CTSO |  |  |
| CG5171 | No ortholog |  |  |
| CG5910 | No ortholog |  |  |
| CG5946 | CYB5R 1-3 |  |  |
| CG6067 | No good ortholog |  |  |
| CG6415 | AMT |  |  |
| CG6484 | SLC2A8 / SLC286 |  |  |
| CG6767 | PRPS1, 2, 1L1 |  |  |
| CG6770 | NUPR1 |  |  |
| CG7322 | DCXR |  |  |
| CG7470 | ALDH18A1 |  |  |
| CG8129 | SRR |  |  |
| CG8586 | No good ortholog |  |  |
| CG9008 | No ortholog |  |  |
| CG9238 | PPP1R3C / 3B |  |  |
| CG9331 | GRHPR |  |  |
| CG9510 | ASL |  |  |

|  |  |
| --- | --- |
| CG9691 | No ortholog |
| CG9886 | GLYCTK |
| CG9928 | No ortholog |
| colt | SLC25A20 |
| Cyp6a23 | TBXAS1 / CYP3A4,5,7,43 |
| Cyp6w1 | CYP3A4, 5, 43 |
| Cyt-b5-r | FADS1, 2, 3 |
| Drat | VAT1L |
| eas | ETNK1, 2 |
| Egfr | EGFR / ERBB2, 3, 4 |
| Est-6 | CES1, 2, 3, 4A, 5A |
| fbp | FBP1, 2 |
| Galt | GALT |
| Gk | GK / GK2, 3P |
| GNBP3 | No ortholog |
| GstE12 | GSTT2B |
| Gyk | GK / GK2, 3P |
| Hn | PAH |
| Idgf4 | CHI3L2 / OGVP1 / CHIT1 /<br>CHI3L1 |
| IM1 | No ortholog |
| IM2 | No ortholog |
| IM3 | SLC25A3 |
| Jheh1 | EPHX1 |
| Mpcp | SLC25A3 |
| MtnA | No good ortholog |
| Npc2g | NPC2 |
| Obp99a | No ortholog |
| Obp99c | No ortholog |
| P5cr-2 | PYCR1, 2, 3 |
| Pepck | PCK1, 2 |
| Pgm | PGM1, 5 |
| ppl | GCSH |
| Prat2 | PPAT |
| Rab9 | RAB9A, B |
| Sodh-1 | SORD |
| Sox102F | SOX5, 6 |
| Spat | AGXT |
| SPE | F10 |
| Spn43Ab | SERPINI1, 2, B1 |
| Tps1 | No ortholog |
| trbl | TRIB1, 2, 3 |
| Tsf1 | MELTF / LTF / TF |
| UGP | UGP2 |
| UK114 | RIDA |
| v | TDO2 |

**Table 5.** CRTC Contra-regulated cardiac genes with conserved CREB binding sites.

| gene | FCA heart cell counts | Human Ortholog | Function | DIOPT |
| --- | --- | --- | --- | --- |
| CG13315 | 466.79 | no ortholog | unknown | - |
| CG9297 | 299.60 | Srl | SR Ca<sup>2+</sup> binding protein | 11 |
| AdipoR | 247.33 | Adipor1 | Glucose/lipid metabolism | 16 |
| CG32549 | 241.77 | NT5C2 | enable 5'-nucleotidase activity. | 15 |
| CG43693 | 240.98 | SLC36A1 | Amino Acid transporter | 13 |
| aralar1 | 229.88 | ARALAR | Mitochondrial Ca<sup>2+</sup> binding transport protein | 18 |
| Prat2 | 198.49 | PRAT | Glutamine amidotransferase | 17 |
| Sodh-1 | 194.95 | SORD | Sorbitol dehydrogenase 1 | 17 |
| CG9331 | 142.51 | GRHPR | glycerate dehydrogenase | 9 |
| trbl | 113.93 | TRIB | Trib family of kinase-like protein | 16 |
| CG3902 | 81.16 | ACADSB | AA transporter | 16 |
| CG5910 | 79.49 | no ortholog | unknown | - |
| Adgf-D | 47.39 | ADGF-C | Adenosine deaminase-related growth factor | 12 |
| Cyp6a23 | 31.55 | CYP3A5 | Cytochrome P450 6a23 | 9 |
| Act42A | 20.81 | ACTB | Actin | 12 |
| **FCA (FlyCellAtlas) myocardial cell count expressing each gene from snRNAseq.**<br>**DIOPT Score (1-19) indicates the number of tools showing homolog between the genes**<br>**(v.9, https://www.flyrnai.org/cgi-bin/DRSC_orthologs.pl)** |  |  |  |  |

**Table 6.**
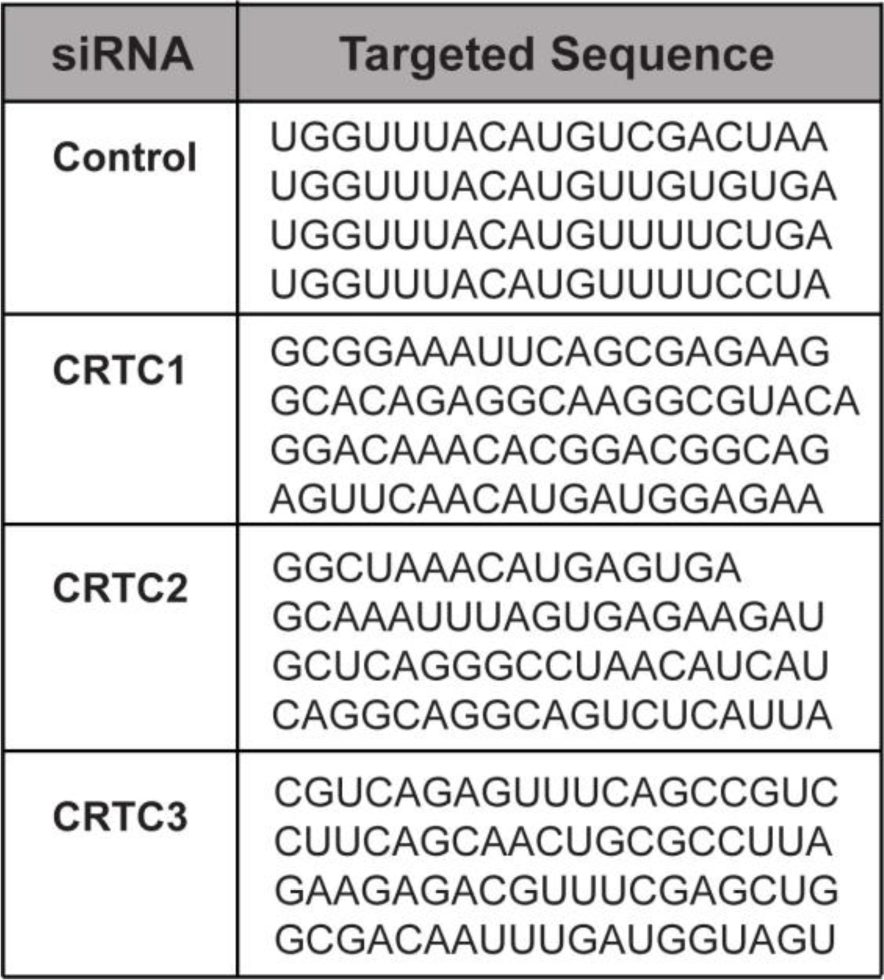
siRNAs targeted sequences for CRTC1, 2 &3. 4 validated sequences for more efficient KD.

**Table 7.** CRTC Expression in hIPS Ventricular-like and Atrial-like Cardiomyocytes.

| Transcript | Gene | Chromosome | Start | Stop | Strand | Length | VCM 1 | ACM 1 | VCM 2 | ACM 2 |
| --- | --- | --- | --- | --- | --- | --- | --- | --- | --- | --- |
| NM_001098482 | CRTC 1 | 19 | 18683615 | 18782334 | + | 6992 | 0.230 | 0.181 | 0.103 | 0.231 |
| NM_015321 | CRTC 1 | 19 | 18683615 | 18782334 | + | 6944 | 0.480 | 0.595 | 0.615 | 0.643 |
| NM_181715 | CRTC 2 | 1 | 1.54E+08 | 1.54E+08 | - | 2678 | 1.788 | 1.677 | 1.548 | 1.574 |
| NM_001042574 | CRTC 3 | 15 | 90529886 | 90645346 | + | 5248 | 1.829 | 1.425 | 1.838 | 1.574 |
| NM_022769 | CRTC 3 | 15 | 90529886 | 90645346 | + | 5251 | 2.519 | 2.477 | 2.082 | 2.572 |

Gene expression in ventricular-like (VCM1) and atrial-like cardiomyocytes (ACM1) at day 12 (early cardiac progenitors) and at day 25 (VCM-2 and ACM-2).

**Table 8.**
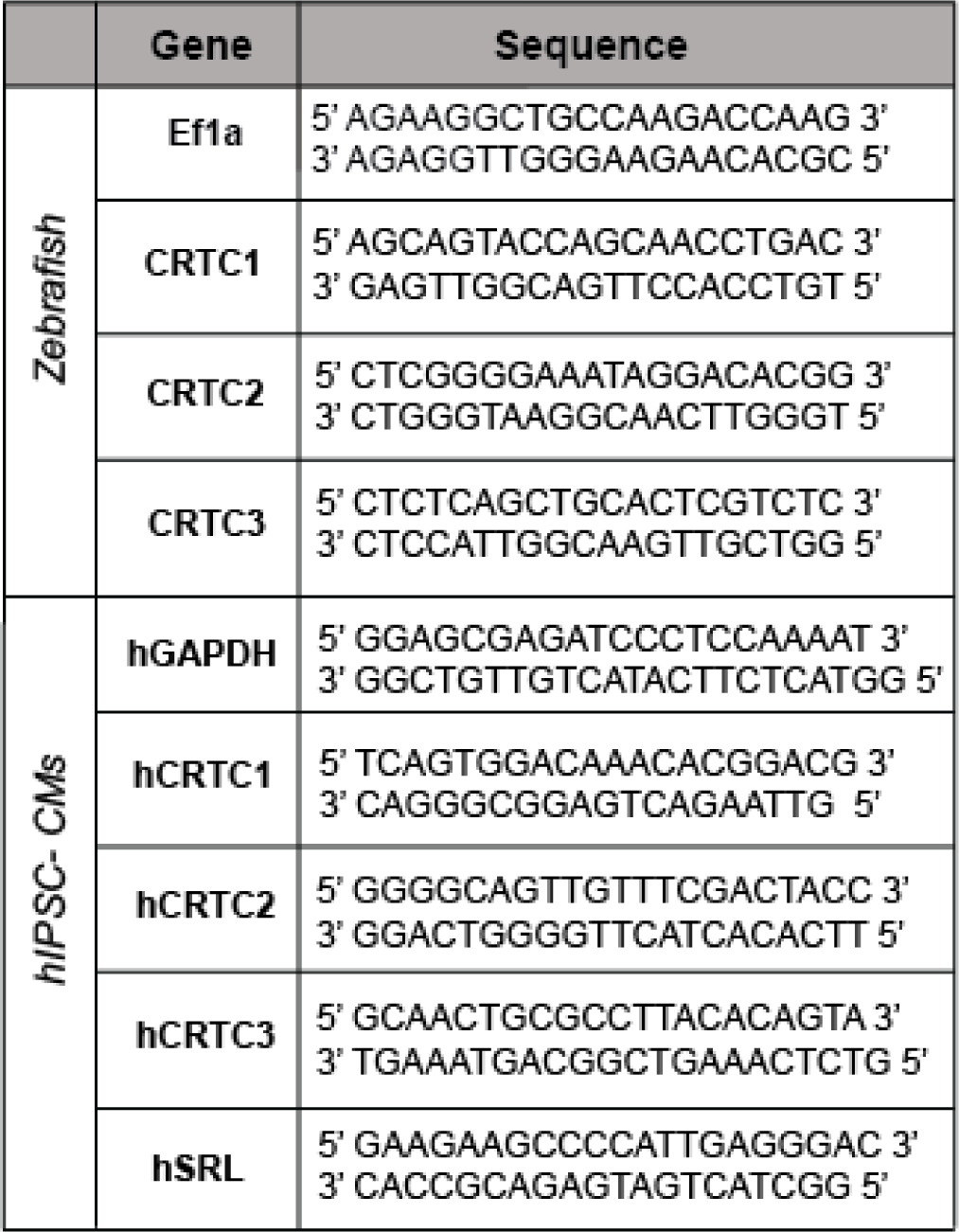
Primers for RT-qPCR of Zebrafish hearts and hIPSC-CMs.

